# Computational design of prefusion-stabilized Herpesvirus gB trimers

**DOI:** 10.1101/2024.10.23.619923

**Authors:** Matthew McCallum, David Veesler

**Affiliations:** Department of Biochemistry, University of Washington, Seattle, Washington, USA; Howard Hughes Medical Institute, Seattle, WA 98195, USA

## Abstract

In the absence of effective vaccines, human-infecting members of the Herpesvirus family cause considerable morbidity and mortality worldwide. Herpesvirus infection relies on receptor engagement by a gH/gL glycoprotein complex which induces large-scale conformational changes of the gB glycoprotein to mediate fusion of the viral and host membranes and infection. The instability of all herpesvirus gBs have hindered biochemical and functional studies, thereby limiting our understanding of the infection mechanisms of these pathogens and preventing vaccine design. Here, we computationally stabilized and structurally characterized the Epstein-Barr virus prefusion gB ectodomain trimer, providing an atomic-level description of this key therapeutic target. We show that this stabilization strategy is broadly applicable to other herpesvirus gB trimers and identified conformational intermediates supporting a previously unanticipated mechanism of gB-mediated fusion. These findings provide a blueprint to develop vaccine candidates for these pathogens with major public health burden.

---

The ubiquitous human-infecting members of the Herpesvirus family cause significant morbidity and mortality globally. Epstein Barr Virus (EBV), a gamma-Herpesvirus, causes 1.3-1.9 % of all human cancers and has been strongly associated with multiple sclerosis^1–3^. Cytomegalovirus (CMV), a beta-Herpesvirus, is a leading cause of hearing loss and permanent disability as a consequence of frequent infection of fetuses^4^. In older adults, the alpha-herpesviruses, Herpes Simplex Viruses (HSV1/2) and Varicella Zoster Virus (VZV), are involved in the etiology of dementia^5–8^. With seroprevalences of >90% for EBV, 45-100% for CMV, ∼50% HSV1, ∼10% HSV2, and 98% for VZV^9–12^, effective countermeasures are urgently needed.

While vaccines are lacking for most Herpesviruses, live-attenuated vaccines exist for VZV which reduce acute disease (Chickenpox) by 87% and reactivation (Shingles) by 51%^13,14,15^. Recently, a subunit vaccine has shown ∼90% efficacy in reducing VZV reactivation^16–18^. This subunit vaccine elicits neutralizing antibodies and cell-mediated immunity against glycoprotein E (gE), which is involved in viral entry^16,17^. Since gE is not conserved or absent in other Herpesviruses, alternative entry glycoproteins have been explored as immunogens for vaccine development. Notably, neutralizing and protective antibodies have been identified against the conserved Herpesvirus glycoprotein H and L, which fold as a heterodimer, (gH/gL) and glycoprotein B _(gB)_^19–28^.

Herpesvirus entry typically begins with gH/gL binding to the host receptor, either directly or with other virus-specific glycoproteins^29^. Upon receptor engagement, gH/gL triggers gB to change conformation and promote fusion of the viral and host membranes^29^. While several gH/gL-targeted neutralizing antibodies compete for binding with host receptors^30^, the mechanisms of most gB-directed neutralizing antibodies remain elusive. The gB ectodomain comprises five subdomains, designated DI, DII, DIII, DIV, and DV, followed by C-terminal transmembrane and cytoplasmic domains^31^ (**Fig 1a-b**). CMV and HSV1 gB share four antigenic sites targeted by neutralizing antibodies, located in DI, DII, DIV, and the flexible N-terminus^32,33,19,32^ with the DIV antigenic site being shared with VZV gB and EBV gB^34–36^. Recently, the DIV-targeted 93k VZV neutralizing antibody was proposed to be incompatible with a DIV-bound conformation of the gB N-terminus, though, the underlying molecular mechanism of neutralization remains unknown^35,37^. Our limited understanding of the mechanisms of action of gB-targeted neutralizing antibody highlights gaps in our knowledge of Herpesvirus fusion.

**Figure 1.**
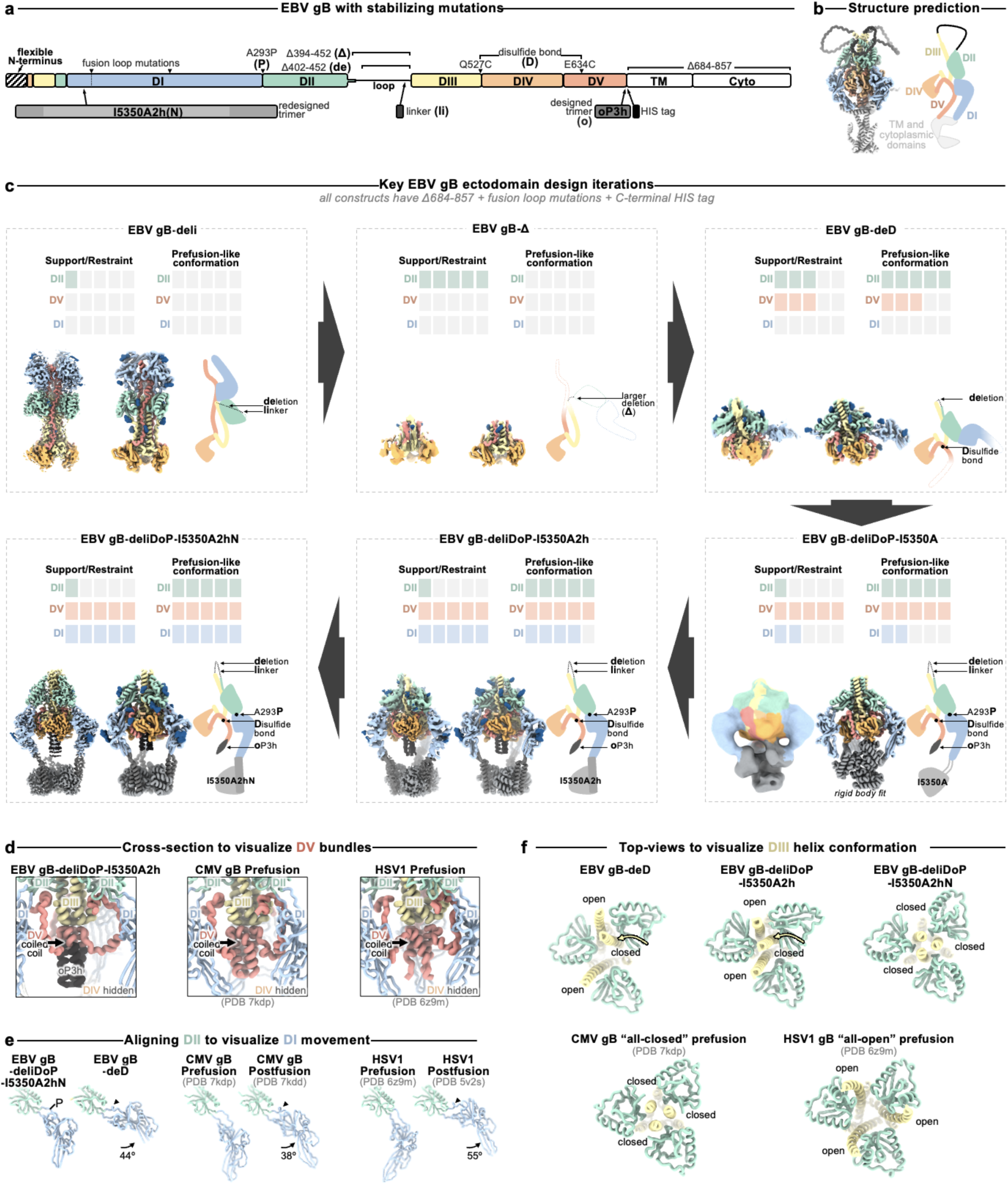
Prefusion-stabilization of the EBV gB ectodomain trimer. **(a)** Schematic of the domain organization of full-length Epstein Barr Virus (EBV) gB with the designed modifications to produce the gB-deliDoP-I5350A2hN ectodomain indicated. All constructs have deletion of the transmembrane and cytoplasmic domains, fusion loop mutations to enhance solubility, and a C-terminally fused polyhistidine tag (**Table S1**). de: deletion of residues 402-452; Δ: deletion of residues 394-452; li: GSPPGSPP loop-linker, D: disulfide bond between residues Q527C and I634C (D); P: A293P mutation; o: oP3H trimer fusion; I5350A2h(N): fusion of the redesigned I5350A. (**b**) Left, ribbon representation of the AlphaFold2-predicted structure of full length prefusion EBV gB using prefusion CMV gB as a template (PDB 7kdp). Right, schematic of the 3D organization of prefusion gB domains in one protomer. (**c**) EBV gB ectodomain design iterations as panels, including gB-deli, gB-Δ, gB-deD, gB-deliDoP-I5350A, gB-deliDoP-I5350A2h, and gB-deliDoP-I5350A2hN. Top of panels, qualitative overview of the relative conformational restraint imposed by mutagenesis, as well as the relative prefusion-like conformation of DII, DV, and DI. Bottom of panels, the locally sharpened CryoEM map (left, surface), model built into the map (middle, ribbons), and schematic of one protomer (cartoon, right) is shown with mutations labeled as in panel (a). A composite map from locally refined and sharpened cryoEM maps is shown for EBV gB-deliDoP-I5350A2h and EBV gB-deliDoP-I5350A2hN. Domain I (DI, blue), domain II (DII, green), domain III (DIII, yellow), domain IV (DIV, orange), domain V (DV, red), glycans (dark blue), oP3h (black), and I5350A2h (gray). (**d**) Comparison of the DV C-termini among EBV gB deliDoP-I5350A2h, CMV gB (PDB 7kdp) and HSV1 gB (PDB 6z9m). (**e**) Comparison of the DI-DII relative orientation and intervening hinge among EBV gB deliDoP-I5350A2hN, EBV gB-deD prefusion and postfusion CMV (PDB 7kdp and 7kdd, respectively), and prefusion and postfusion HSV1 gB(PDB 6z9m and 5v2s, respectively). Black triangles show α-helically restructured hinge residues characteristic of the postfusion state; P: A293P mutation (**f**) Comparison of the organization of the DIII central helices among EBV gB-deD, EBV gB-deliDoP-I5350A2h, EBV gB-deliDoP-I5350A2hN, prefusion CMV gB (PDB 7kdp), and prefusion HSV1 gB (PDB 6z9m).

No structural information exists for prefusion EBV gB or any other gamma-Herpesviruses, hindering our ability to design countermeasures against this important class of pathogens. Prefusion HSV1 gB H516P as well as the corresponding VZV gB mutant were visualized at ∼1 nm resolution using cryoET, revealing their domain organization^38,39^. Furthermore, a cryoEM structure of full-length CMV gB (stabilized with a CMV-specific inhibitor and covalent crosslinking) was recently described^40^. Comparing prefusion and postfusion conformations of these gB structures show that DII linked to DI (which harbors hydrophobic fusion loops) and DV (anchored to the viral membrane) are reoriented ∼180° during the membrane fusion reaction relative to DIII and DIV^41^. For class III fusion proteins, such as Herpesvirus gB, the fusion loops are thought to reorient for insertion into the host membrane before structural rearrangements of the domain directly anchored to the viral membrane^42–44^.

Although viral fusion glycoproteins are finely tuned to promote entry with exquisite spatial and temporal control, irreversible refolding to the postfusion state often occurs upon purification. Therefore, stabilizing them in the prefusion conformation is typically necessary to produce an immunogen eliciting potent immune responses^45–47^. For example, the 2P and DS-CaV1 prefusion-stabilizing mutations enhance glycoprotein stability and enable elicitation of potent immune responses for approved vaccines against SARS-CoV-2 and RSV, respectively^47–49^.

Whereas SARS-CoV-2 and RSV use class I fusion proteins, Herpesvirus gB is a homotrimeric class III fusion protein, which is markedly distinct and much less well characterized with limited prefusion-stabilization success despite extensive prior attempts^50,51^. The H516P HSV1 gB mutation increases the prefusion to postfusion ratio in HSV1 gB, as does the corresponding mutation in VZV gB. However, the applicability of this strategy is limited to full-length membrane-embedded protein and fails to stabilize the prefusion recombinant ectodomain trimer adequately^38^. No prefusion-stabilizing mutations are known for soluble gB ectodomains of any Herpesvirus; consequently, there are no isolated prefusion gB-specific neutralizing antibodies. Accordingly, stabilizing the prefusion gB conformation would not only further our understanding of herpesvirus-mediated membrane fusion but also facilitate the development of countermeasures against this important family of pathogens.

Here, we stabilized the prefusion EBV gB ectodomain trimer using recent advances in structure-based and machine-learning guided protein design approaches, and identified conformational intermediates that support an alternative mechanism of gB-mediated fusion. We show that the stabilization strategy is broadly applicable to alpha-, beta-, and gamma-Herpesviruses, pointing to a shared fusion mechanism and paving the way to develop vaccine candidates against these pathogens.

## Results

### Preventing Domain II reorientation in gB

We hypothesized that deleting the loop between DII and DIII would restrict DII—and consequently gB—to the prefusion conformation, as the loop spans a distance of 20 Å in the predicted prefusion conformation and 40 Å in the postfusion conformation (**Fig 1a-b** and **Extended Data Fig. 1a-b**). This loop is not anticipated to play a significant role in fusion, aside from the presence of a polybasic furin cleavage site which enhances (but is not necessary for) gB-mediated fusion^52^. Therefore, we deleted this 51-residue loop between DII and DIII (residues 402-452) and replaced it with an eight residue linker (herein gB-deli). CryoEM structure determination unexpectedly revealed that gB-deli adopted the postfusion conformation (**Fig 1c**, **Extended Data Fig. 2a**, and **Table S2**), underscoring the need for further stabilization.

We subsequently designed gB-Δ, with a deletion of the entire DII-DIII loop along with that of the adjacent residues 394-401 (predicted to form an ⍺-helix bridging DII to the loop), which prevented refolding to the postfusion conformation (**Extended Data Fig. 3a-c**). However, DI and DII adopted random orientations relative to the rest of the ectodomain, inconsistent with the prefusion conformation (**Extended Data Fig. 3c**), and were not resolved upon cryoEM analysis (**Fig 1c**, **Extended Data Fig. 2b**, and **Table S2**). The gB-Δ structure reveals that DV interacts with DIII in a postfusion-like manner, thereby occluding the site on DIII that would be occupied by DII in the prefusion state (**Extended Data Fig. 1c**). This observation rationalizes the conformational heterogeneity of DI and DII, as DII cannot bind to its prefusion binding site on DIII, while the 394-401 loop deletion prevents DII from moving to its postfusion binding site on DIII.

### Preventing Domain V flipping in gB

Since DV moves to the postfusion conformation in gB-Δ, we targeted this domain to introduce additional mutations for stabilization of the prefusion state. Using ProteinMPNN^53^ along with the AlphaFold2-predicted^54^ structure of prefusion EBV gB, we identified and screened ∼100 possible stabilizing mutations. An ectodomain that adopts the postfusion conformation (gB with the 402-452 residue deletion, gB-de) was used to enable screening of mutations by 2D classification of negatively stained samples. We identified an intermolecular disulfide bond between DIII and DV (Q527C-E634C) that prevents the transition to the postfusion conformation of the gB-de ectodomain (**Extended Data Fig. 3d**). Non-reducing SDS-PAGE analysis of this ectodomain construct (gB-deD herein) was consistent with formation of the designed intermolecular disulfide bond (**Extended Data Fig. 3d**).

To probe the role of the loop deletion in stabilization, we reintroduced an eight residue loop-linker (herein gB-deliD) which resulted in dampened intermolecular disulfide bond formation and refolding to the postfusion state (**Extended Data Fig. 3e**). This indicates that the conformational restriction of DII imposed by the loop deletion contributed to enhancing the stability of prefusion gB-deD, and that release of this restriction by reintroducing the loop-linker in gB-deliD favors the postfusion state. CryoEM structure determination of gB-deD revealed prefusion-like interfaces between DII and DIII as well as between DIV and DV, whereas the predicted DV C-terminal 3-helix coiled-coil was not resolved in the map (**Fig 1c**, **Extended Data Fig. 2c**, and **Table S2**). In summary, we identified the Q527C-E634C disulfide bond which participates in stabilizing the prefusion conformation of gB DII and DV through interprotomer crosslinking.

### A designed helical trimer stabilizing Domain V

To further enhance the stability of DV, we computationally designed a seven residue trimerization motif, designated oP3h, to promote the formation of the C-terminal bundle (**Fig 2a**). To evaluate the ability of oP3h to stabilize well-characterized fusion proteins, we replaced the C-terminal foldon trimer of the SARS-CoV-2 S-2P ectodomain^55^ with oP3h motifs with varying number of repeats (**Fig 2b**). Recombinant production yield was inversely correlated with the number of oP3h repeats with three repeats providing optimal balance for prefusion stabilization of S-2P trimers (**Fig 2b** and **Extended Data Fig. 4a-h**). CryoEM structure determination of the SARS-CoV-2 S-2P ectodomain with oP3h (three RINEIER or four RINAIET repeats) confirmed the predicted structure of oP3h and the accuracy of our designs (**Fig 2c**, **Extended Data Fig. 4i-j**, and **Table S3**).

**Figure 2.**
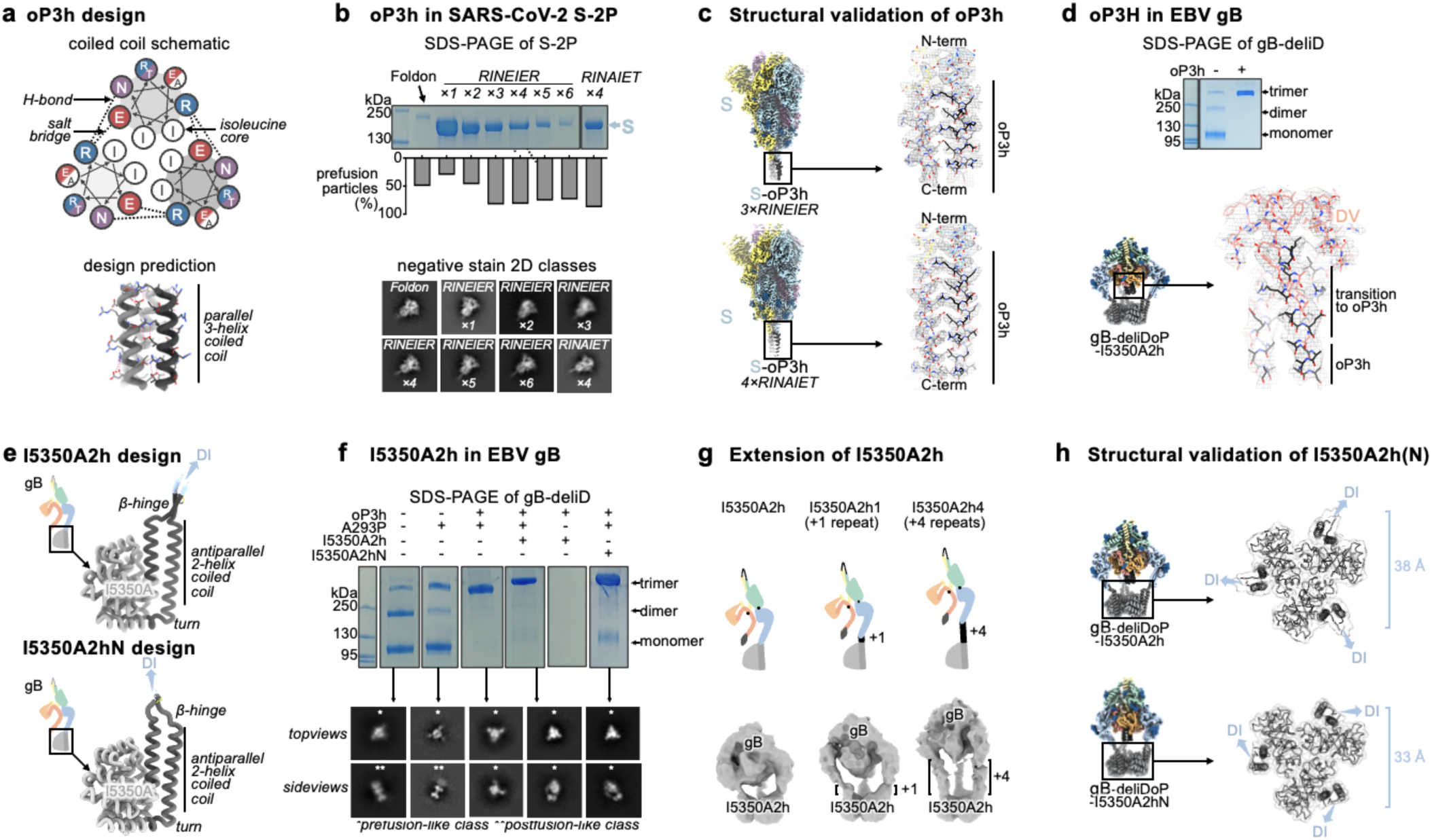
Design and validation of oP3h and I5350A2hN trimers for prefusion gB stabilization. **(a)** Wheel diagram (top) and AlphaFold2-predicted structure of the oP3h trimer, shown in ribbon representation with residues labeled (bottom). **(b)** Top, Coomassie stained SDS-PAGE of SARS-CoV-2 S ectodomains with C-terminally fused foldon, one to six oP3h (RINEIER) repeats, or with four oP3h (RINAIET) repeats. Glycoproteins were expressed and purified in parallel and normalized to a standard volume for comparison of relative yields after affinity purification. Bottom, the proportion of S ectodomains (after affinity purification) that are clearly folded in the prefusion state, as determined by negative stain 2D classification. Exemplar prefusion 2D class averages are shown underneath. (**c**) CryoEM maps of SARS-CoV-2 S-oP3h with three RINEIER (top) or four RINAIET (bottom) C-terminal repeats with each protomer colored distinctly. Insets show local refinement maps (mesh) and refined models (ribbons) encompassing the oP3h fusion. **(d)** Top, Coomassie stained non-reducing SDS-PAGE analysis of EBV gB deliD with (+) and without (-) oP3h. Bottom, cryoEM map (mesh) and model (ribbons) of the EBV gB deliDoP-I5350A2h domain V (DV) and oP3h fusion. **(e)** Design and AlphaFold2-predicted models of I5350A2h and I5350A2hN, showing the predicted relative orientation of DI to which they are fused. **(f)** Left, Coomassie stained non-reducing SDS-PAGE and 2D EM class averages of negatively stained gB-deliD ectodomains with (+) and without (-) oP3h, A293P (P), or I5350A2h. (**g**) Top, Schematic showing the addition of 1 (center) or 4 (right) heptad repeats to the coiled coils of I5350A2h. Bottom, EM reconstructions of negatively stained EBV gB deliDoP-I5350A2h with heptad repeats added to the coiled coils of I5350A2h. (**h**) Local refinement cryoEM maps (transparent surface) and model (black ribbons) of the I5350A2h or I5350A2hN regions within EBV gB deliDoP-I5350A2h and EBV gB deliDoP-I5350A2hN, respectively; blue arrows, the relative orientation of DI. The diameter at which DI is held by I5350A2h or I5350A2hN is labeled on the right.

We subsequently fused oP3h in-frame with the EBV gB DV C-terminus using the gB-deliD ectodomain, including substitutions that gradually transition DV to oP3h (herein gB-deliDo) (**Fig 2d**). oP3h fusion effectively promoted formation of the Q527C-E634C intermolecular disulfide bond, as observed by non-reducing SDS-PAGE, consistent with DV adopting the prefusion conformation (**Fig 2d**). Given that gB-deliDo does not transition to the postfusion conformation (**Extended Data Fig. 3f**), increased DV stabilization compensates for the enhanced conformational freedom of DII resulting from the presence of the flexible loop-linker region.

### Domain I hinge stabilization in gB

While gB-deD and gB-deliDo resist DV and DII refolding to the postfusion state, the cryoEM structure of gB-deD and the negative stain reconstruction of gB-deliDo feature DI pointing laterally away from the gB ectodomain, unlike the DI clamping of DV observed for prefusion CMV gB and HSV1 gB (**Fig 1c**, **Extended Data Fig. 3g, and Extended Data Fig. 5**).

Comparing the relative orientation of DI and DII in gB-deD with that of prefusion and postfusion CMV gB shows that DI has a postfusion-like orientation in which residues 290-295 adopt an ⍺-helical conformation (**Extended Data Fig. 5f-i**). Analogous domain rotations between prefusion and postfusion states, including the ⍺-helical hinge restructuring, are conserved in distantly related Rhabdovirus class III fusion proteins^56^ (**Extended Data Fig. 5j-k**). A similar DI orientation and DI hinge structure are observed in a recent preprint describing disulfide stapling of CMB gB DV and DI.; This DI-DII relative orientation along with the absence of DV C-terminal bundling suggest a similar degree of stabilization to EBV gB-deD^57^. We hypothesized that the postfusion DI hinge is rigidly locked, whereas the extended conformation observed for the prefusion DI hinge enables rotation – consistent with a highly stable postfusion conformation and metastable prefusion conformation. Using ProteinMPNN^53^, the A293P mutation was selected to stabilize the prefusion-like DI hinge conformation. Consistent with decreased rigidity of the hinge, negative stain 2D classification and 3D reconstructions of gB-deliD or gB-deliDo harboring A293P (gB-deliDP and gB-deliDoP, respectively) show that DI is flexible in the prefusion and postfusion conformations (**Fig 2f** and **Extended Data Fig. 3h**). Just as DII and DV stabilization (gB-deliDo) does not prevent DI reorientation, the A293P substitution does not suppress large scale DII postfusion-like conformational changes (observed via negative staining EM), nor increased prefusion DV stability (observed via non-reducing SDS-PAGE analysis of Q527C-E634C bond formation) (**Extended Data Fig. 3e-h**). These data demonstrate that DI hinge conformational changes occur independently of DII and DV conformational changes in the gB ectodomain, defining DI rotation as a third fusogenic conformational change, separate from DII and DV flipping.

### A redesigned trimeric fusion enhances gB stabilization

In the context of full length prefusion gB, the fusion loops are anticipated to be embedded in the viral envelope limiting the range of DI motion. To simulate the envelope-bound orientation of DI, we genetically inserted the I5350A trimeric fusion domain^58^ in the DI fusion loops using flexible linkers. CryoEM analysis of gB-deliDoP-I5350A revealed that the flexible linkers allowed I5350A to rotate markedly relative to gB (**Fig 1c**, **Extended Data Fig. 2d**, **Extended Data Fig. 3j**, and **Table S2**). To prevent this, I5350A was redesigned to harbor a structured and extendable link to DI (designated I5350A2h), and evaluated with the gB-deliDo and gB-deliDoP constructs (**Fig 2e-h**, and **Extended Data Fig. 3k-m**). In the absence of the A293P mutation, we could not detect production of the gB-deliDo ectodomain fused to I5350A2h (**Fig 2f**), consistent with a rigid postfusion-like DI hinge positioning DI in a manner incompatible with the presence of the I5350A2h trimer. Conversely, in the presence of the A293P mutation, the EBV gB trimer folded as designed, enabling visualization of I5350A2h and of oP3h in the cryoEM structure of EBV gB-deliDoP-I5350A2h (**Fig 2d**, **Fig 2h**, **Extended Data Fig. 3k**, **Extended Data Fig. 6a** and **Table S2**) In the structure of EBV gB-deliDoP-I5350A2h, the diameter of I5350A2h positions DI too far apart for all DI domains to directly contact DIV and DV (**Extended Data Fig 5l-o**). As this interaction is observed in prefusion CMV gB and HSV1 gB, we removed four residues from the redesigned I5350A2h helices, thereby bringing DI closer to the rest of the trimer and reorienting the subsequent beta strands by ∼30°. This trimeric fusion was designated I5350A2hN and enabled visualization of DI contacting DV and DIV in the cryoEM structure of EBV gB-deliDoP-I5350A2hN (**Fig 2e-h**, **Extended Data Fig. 3n**, **Extended Data Fig 5l-o**, **Extended Data Fig. 6b**, and **Table S2**).

### The prefusion stabilized EBV gB trimer

All key structural features in the EBV gB-deliDoP-I5350A2hN ectodomain matched the predicted prefusion EBV gB architecture (**Fig 1c-e**, **Extended Data Fig. 1**, and **Extended Data Fig. 5**). Comparing the structure of EBV gB-deliDoP-I5350A2hN to prefusion CMV gB and HSV1 gB structures reveals many similarities including the orientation of DII and the conformation of DV (**Fig 1c,d**). Furthermore, a 44° rotation of DI relative to DII occurs when comparing the EBV gB-deliDoP-I5350A2hN and gB-deD structures, concurring with CMV and HSV1 gB prefusion and postfusion structures, which enables DI to clamp DV in prefusion gB (**Fig 1d,e**). The central helices of DIII from EBV gB-deliDoP-I5350A2hN are all tightly packed or closed coiled coils, as they are in the CMV gB prefusion structure^40^, which is distinct from the HSV1 gB prefusion structure where these helices are splayed open^38^(**Fig 1f**). This contrasts with the central helices of DIII from EBV gB-deD and EBV gB-deliDoP-I5350A2h, which adopt an intermediate state with two open and one closed DIII helix (**Fig 1f**), supporting a role for DI positioning in modulating the conformation of DIII. Overall, EBV gB-deliDoP-I5350A2hN is structurally similar to full length prefusion CMV gB and HSV1 gB (5.2 Å and 5.8 Å RMSD^Cɑ^, respectively, compared to 6.1 Å RMSD^Cɑ^ for CMV gB vs HSV1 gB). The designed stabilization strategy enables recapitulating the key structural features anticipated for prefusion gB, validating this design, and creating a template for prefusion-stabilization of gB from related viruses.

### A broadly applicable stabilization strategy

Given the local structural conservation of the mutated regions in EBV gB-deliDoP-I5350A2h(N) with CMV gB and HSV1 gB, we hypothesized that this stabilization strategy would be applicable to other human-infecting Herpesviruses. Accordingly, the CMV, HHV6B, and HSV1 gB-deliDoP-I5350A2h ectodomain trimers appear similar to that of EBV upon negative staining and 2D classification of EM data (**Extended Data Fig. 7a,c** and **Extended Data Fig. 8a,c**). We further found that a subset of these mutations—gB-deliD—were sufficient for prefusion-stabilization of the HSV1 gB ectodomain and partial stabilization of the VZV gB ectodomain (**Extended Data Fig. 8a,c**). The L687K mutation in DV was introduced into VZV gB-deliD to further hinder transition to the postfusion conformation through predicted charge repulsion effects expected to solely occur in the postfusion state (**Extended Data Fig. 1b** and **Extended Data Fig. 8c**). HSV1 gB-deliD and VZV gB deliD formed dimers of prefusion trimers with their fusion loops oriented towards each other (**Extended Data Fig. 8a,c** and **Fig 3g**). CryoEM analysis of HSV1 gB-deliD, VZV gB-deliD + L689K, CMV gB-deliDoP-I5350A2h, HHV6B gB-deliDoP-I5350A2h confirmed folding in the prefusion conformation, as evidenced by the orientations of DI, DII, and DV (**Fig. 3**, **Extended Data Fig. 7, Extended Data Fig. 8**, and **Table S4**). Collectively, these data show that the prefusion-stabilizing mutations designed here are broadly applicable to divergent Herpesvirus gBs from multiple subfamilies.

**Figure 3.**
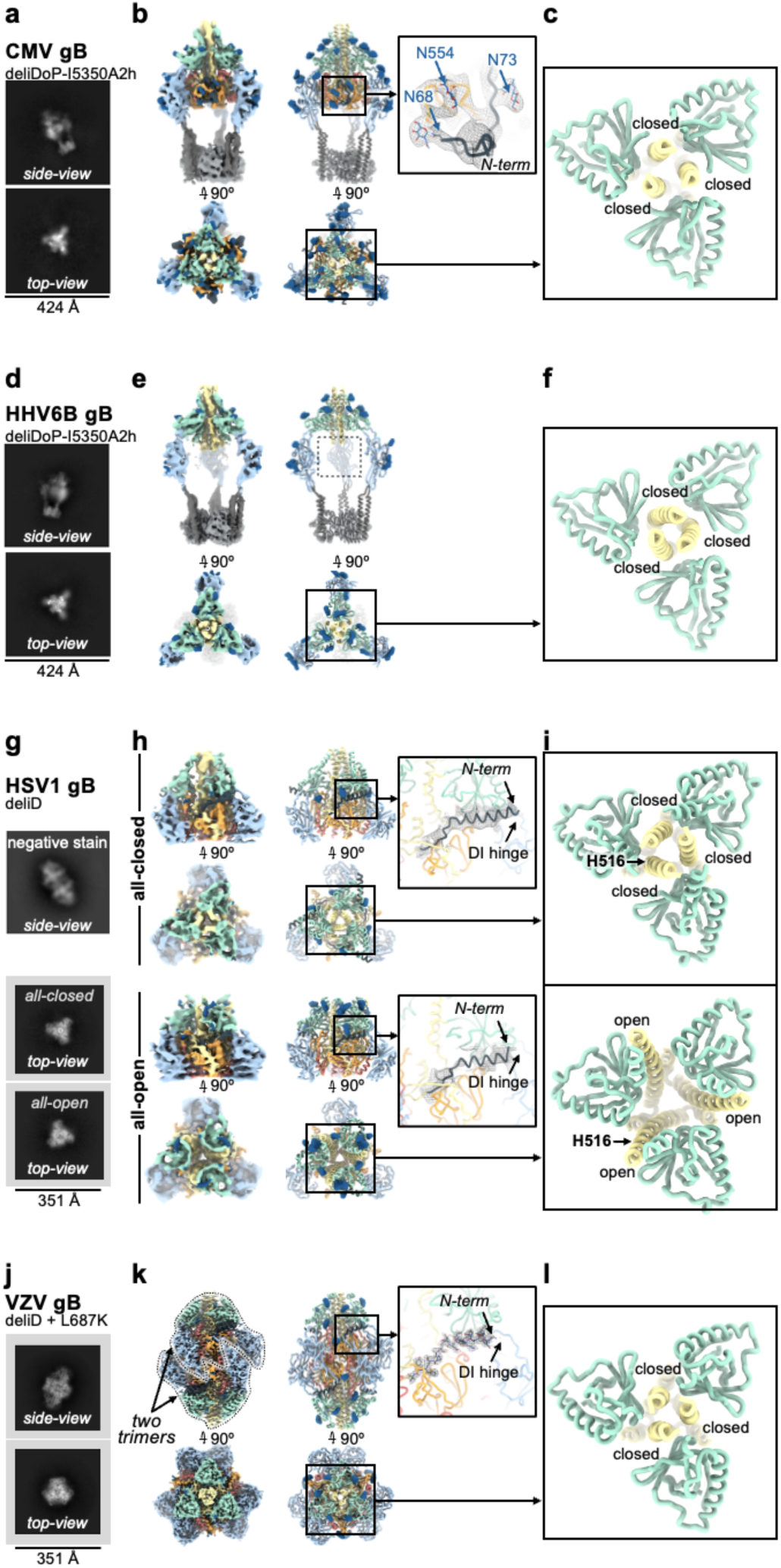
A broadly generalizable herpesvirus gB prefusion-stabilization strategy. Structural analysis of CMV gB-deliDoP-I5350A2h (a-c), HHV6B gB-deliDoP-I5350A2h (d-f), HSV1 gB-deliD (g-i), and VZV gB-deliD + L687K (j-l). (a,d,g,j) Selected cryoEM 2D classes representing side- and top-views. For HSV1 gB-deliD (**g**) preferred particle orientation limited cryoEM 2D classes to top-views (bottom two panels), so the side-view from negative stain is also shown (top). (**b,e,h,k**) CryoEM maps (left) and ribbon diagrams of the corresponding model (right) with orthogonal views. The composite map of locally refined and sharpened cryoEM maps are shown for CMV and HHV6B-gB-deliDoP-I5350A2h (**b,e**), the locally refined and sharpened cryoEM maps are shown for HSV1 gB-deliD (**h**), and the sharpened cryoEM map is shown for VZV gB-deliD + L687K (**k**). Insets (right) show zoomed-in views of the N-terminal residues (black). (**c,f,i,l**) View of domain III (DIII, yellow) central helices and domain II (DII, green). Individual chains are labeled as closed or open by comparison with Fig 1f. The position of residue H516 is labeled for HSV1 gB-deliD.

### Domain III occupies distinct conformational states

The most obvious distinction between Herpesvirus prefusion gB ectodomains is the conformation of the DIII helices. While EBV gB-deliDoP-I5350A2h displays a two-closed/one-open DIII conformation, EBV gB-deliDoP-I5350A2hN, CMV gB-deliDoP-I5350A2h, HHV6B gB-deliDoP-I5350A2h, HSV1 gB-deliD, and VZV gB-deliD + L689K ectodomains adopt an all-closed DIII conformation (**Fig 3c,f,l**), similar to the prefusion structure of full length CMV gB (**Fig 1f**). HSV1 gB-deliD was also identified in a second conformation with all-open DIII helices (**Fig 3h,i**), similar to the structure of full length prefusion HSV1 gB^38^ (**Fig 1f**). The observation that the CMV gB-deliDoP-I5350A2h and HSV1 gB-deliD ectodomains faithfully recapitulate the DIII conformations observed in the corresponding full length glycoprotein structures is indicative of native folding of these constructs.

### Resolving the N-terminal antigenic site

Density consistent with the previously unresolved N-terminal residues of gB from CMV, HSV1, and VZV was present in our cryoEM maps (**Fig 3b,h,k**). The CMV gB and HSV1 gB N-terminal residues are targeted by neutralizing antibodies and these residues are known to play a functional role in VZV gB-mediated membrane fusion^32,37,59,60^. For CMV gB-deliDoP-I5350A2h, this additional density was consistent with the predicted structure^54^ of residues 67-86, bound to DIV, and further supported by visualization of the N68 and N73 glycans in the cryoEM map (**Fig 3b**). For HSV1 gB-deliD, this additional density was consistent with the predicted^54^ helical structure of residues 86-110, bound to the DI hinge (**Fig 3h**). In the low pH and postfusion HSV1 gB structure, these residues adopt a markedly distinct conformation bound to DIV^48^, similar to the DIV-bound N-terminal residues of CMV gB-deliDoP-I5350A2h (**Extended Data Fig. 9a**). For VZV gB-deliD, the N-terminal region was resolved at high-enough resolution to enable de novo model building of residues 100-116, adopting a helical conformation and interacting with the DI hinge (**Fig 3k**). Numerous contacts between the VZV gB N-terminal helix and the DI hinge region support a role in stabilizing the prefusion hinge conformation (**Fig 3k and Extended Data Fig. 9a,b**), and the corresponding residue pairs remain complementary in HSV1 gB (**Extended Data Fig. 9b**). Mutation of the N-terminal VZV gB residues involved in this interaction (^109^KSQD^112^ to ^109^AAAA^112^) disrupt viral propagation^37^, underscoring the functional relevance of these contacts.

### gB-targeted neutralizing antibody mechanisms

To provide insights into the mechanism of inhibition of gB-targeted neutralizing antibodies, we aligned available structures of antibody-bound postfusion gB to the prefusion gB structures described here. Comparing the prefusion and postfusion conformations of the N-terminal antigenic site, the 93k VZV neutralizing antibody^35^ would selectively prevent N-terminus binding to DIV but not to the DI hinge (**Extended Data Fig. 9c**). The 3A5 EBV neutralizing antibody^34^ epitope overlaps with 93k and might function similarly (**Extended Data Fig. 9c**). Conversely, binding, restructuring, and sequestering of the N-terminal antigenic site by TRL345-like antibodies^60–62^ would prevent binding of the N-terminus to DIV or possible binding to the DI hinge^60–62^(**Extended Data Fig. 9c**). The CMV neutralizing antibodies, SM5^63^ and 1G2^32^, surround and contact the DI hinge and may thus block binding of the N-terminus or directly manipulate hinge conformation (**Extended Data Fig. 9c**). While EBV neutralizing antibody 3A3^34^ and HSV1 neutralizing antibody D48^36^ target DII, near the SM5 epitope, they do not approach the DI hinge (**Extended Data Fig. 9c**). Alternatively, it is expected that 3A3 would promote the all-closed conformation of gB by clashing with two adjacent open gB protomers (**Extended Data Fig. 9c**), while the D48 antibody would restrict the motion of the ⍺-helix between DII and the loop to DIII, impeding DII flipping (**Extended Data Fig. 9c**). This analysis supports that antibodies that neutralize Herpesviruses dysregulate key fusogenic conformational changes, including DIII closing, DI hinge restructuring, and DII flipping.

### Inference of the gB-mediated fusion mechanism

The iterative process of gB prefusion-stabilization provided key insights into the mechanism of gB-mediated membrane fusion (**Fig 4**). For class III fusion proteins like gB, it has been presumed that the fusion loops reorient for insertion into the host membrane prior to structural rearrangements of the domain directly anchored to the viral membrane^42,43^. This would imply that DII rotates first, followed by DV (**Fig 4a-c**). However, our data show that DV can transition to a postfusion conformation even if DII does not (see gB-Δ, **Fig 1c**). In contrast, increased DV stability prevents DII from flipping, even with minimal DII restraint (see gB-deliDo, **Extended Data Fig. 3e-f**). These data suggest that DV reaches its postfusion conformation prior to DII during membrane fusion, concurring with the fact that DI (linked to DII) surrounds DV in all Herpesvirus postfusion gB structures (and distantly related Baculovirus and Thogotovirus class III fusion proteins) (**Fig 4f** and **Extended Data Fig. 10**). Therefore, EBV gB-Δ likely represents a conformational intermediate of the gB fusion reaction (**Fig 4f**).

**Figure 4.**
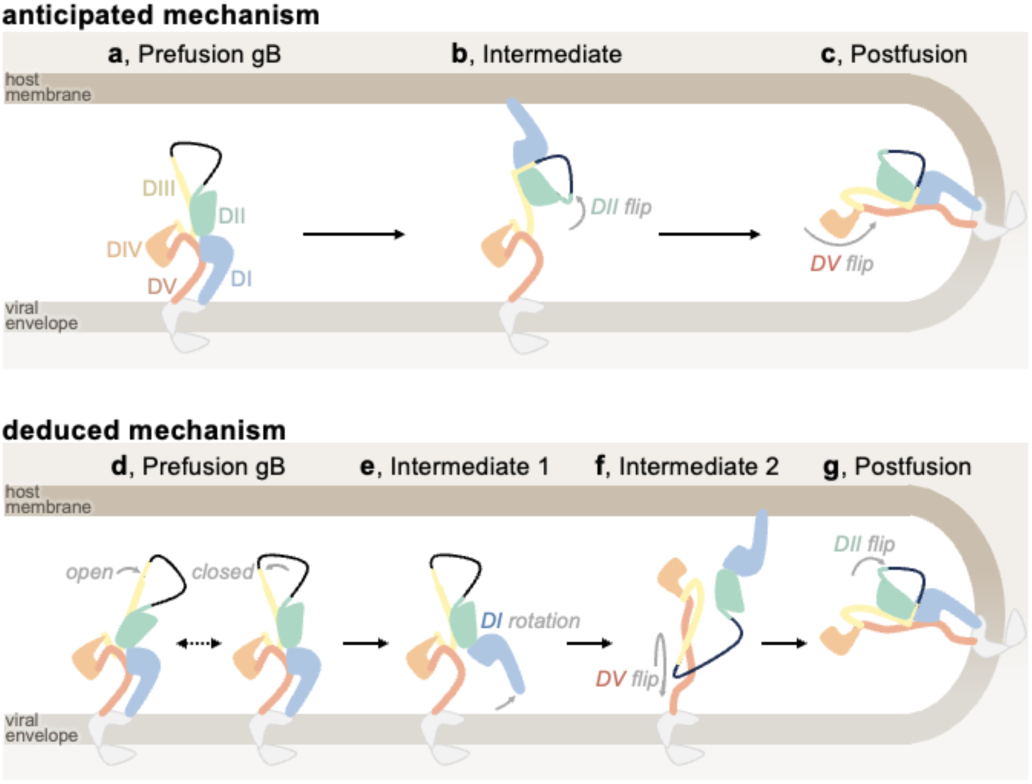
Anticipated and deduced molecular mechanisms of gB fusion. Domain I (DI, blue), domain II (DII, green), domain III (DIII, yellow), domain IV (DIV, orange), and domain V (DV, red) are shown, in addition to the loop between DII and DIII (black line), cytoplasmic and transmembrane portions of gB (grey), the viral envelope (light brown), and host membrane (dark brown). (**a-c**) Cartoon schematic of the anticipated mechanism of gB-mediated fusion at the outset of the study: starting from prefusion gB (**a**), DII flips about DIII, positioning DI to bind the host membrane, (**b**), and then DV flips about DIII, yielding the postfusion conformation while merging viral and host membranes (**c**). (**d-g**) The mechanism of gB-mediated fusion which is consistent with the analysis of gB ectodomains herein: prefusion gB DIII may exist in open and/or closed conformations, likely depending on the virus (**d**). The hinge between DI and DII restructures leading to DI rotation; thereby pulling the fusion loops from the viral envelope while releasing DV (**e**). DV refolds to its postfusion conformation and replaces DII on DIII, releasing DII. Since DV is anchored to the viral envelope, this conformational change reorients the entire ectodomain and positions the DI fusion loops for binding the host membrane (**f**). DII binds its postfusion interface on DIII, yielding the postfusion gB conformation (**g**).

Given that DV is surrounded by DI in both prefusion and postfusion states, (**Extended Data Fig. 5**), conformational changes leading to membrane fusion likely involve DI releasing prefusion DV. A DI conformation that releases its clamp on DV was observed in gB-deD and gB-deliDo ectodomains, with DI pointing laterally away from gB (**Fig 1c**, **Extended Data Fig. 3g**, and **FigS6**). Furthermore, DI reorientation would pull the fusion loops out of the viral envelope—a necessary step for fusion (**Fig 4e**). gB-deliDo may thus represent another conformational intermediate of the gB fusion reaction (**Fig 1e** and **Fig 4e**). Conformational changes induced by DI reorientation are not limited to the release of DV, as the DIII conformation is also changed in EBV gB-deliDoP-I5350A2hN relative to EBV gB-deliDoP-I5350A2h.

Across prefusion-stabilized gB ectodomains, we observed the central DIII helices in open or closed conformations (**Fig 1f** and **Fig 3c,f,i,l**). As HSV1 gB-deliD can adopt all-open and all-closed conformations, these structural rearrangements may be reversible for some viruses (**Fig 3h-i** and **Fig 4d**). In HSV1 gB, the H516P mutation localizes to the central DIII helices (**Fig 3g**), where a proline substitution is anticipated to disrupt the helical secondary structure and DIII bundling^39^. Consistent with this possibility, the all-closed conformation was not observed for full length HSV1 gB with H516P, nor in VZV gB with the corresponding mutation^38,39^. Incompatibility with the all-closed conformation would explain the ability for this proline substitution to provide partial stabilization of HSV1 gB and VZV gB in the all-open prefusion state if the all-closed prefusion conformation was necessary for subsequent conformational changes. Indeed, postfusion DV binds along all-closed DIII helices implying that DV rearrangement depends on the all-closed conformation (**Extended Data Fig. 1b**).

Overall, these findings support an alternate model of gB fusion in which prefusion gB may exist with open and closed conformations of the central helices (**Fig 4d**). DI hinge restructuring would rotate DI and release DV, while dislodging the DI fusion loops from the viral envelope (**Fig 4e**). Once the gB DIII helices are all-closed and DI has rotated, DV could flip to the post-fusion conformation, which would orient DI fusion loops for binding the host-membrane and dislodge the prefusion bound conformation of DII from DIII (**Fig 4f**). Finally, reassociation of DII with the rest of the gB ectodomain in a postfusion conformation would pinch the viral and host membranes and promote fusion (**Fig 4g**).

## Discussion

We leveraged machine-learning- and structure-guided approaches to stabilize the prefusion conformation of the EBV gB ectodomain, paving the way for its evaluation as a vaccine candidate. During this process, we stabilized conformational intermediates of EBV gB, suggesting a plausible mechanism of gB-mediated fusion that may extend to all class III viral fusion proteins. Additionally, we designed trimeric proteins to stabilize the metastable prefusion gB, demonstrating applicability to other fusion glycoproteins, as shown with SARS-CoV-2.

Finally, we applied the designed prefusion-stabilizing mutations to CMV, HHV6B, HSV1, and VZV gBs, showcasing the generalizability of our approach and revealing conserved architectural principles among these viruses. The tools developed here provide a molecular blueprint for designing next generation Herpesvirus vaccines and antivirals to address major unresolved public health challenges.

The conformational snapshots obtained suggest an alternative model of gB-mediated fusion involving movement of the central helices to the all-closed conformation and rotation of DI, followed by DV and subsequent DII rotation. Multiple checkpoints are a common theme across viral entry proteins, as they must exhibit exquisite control in the timing and specificity of the irreversible fusogenic conformational changes they undergo^64^. The open and closed gB conformations described here are reminiscent of those for the HIV-1 Env class I fusion protein, which exhibits similar structural rearrangements to host-receptor bound entry intermediates^65–68^.

Reminiscent of the rotation of gB DI, domain rotation exposing the fusion loops is observed in the class II fusion proteins of Flaviviruses, representing an immunogenic fusion intermediate triggered by low pH^69–75^. By analogy, these gB conformational changes may correspond to critical fusogenic triggers.

Indeed, there are clues that DI hinge restructuring could be a key player in fusogenic triggering. While we show that the N-terminus of VZV and HSV1 gB is bound to the DI hinge, an alternate DIV-bound conformation was captured for HSV1 gB at low pH, which is a fusion trigger for this virus^76,77^. Therefore, we hypothesize that for at least alpha-Herpesviruses the N-terminus maintains the DI hinge in its prefusion state until fusogenic triggering. Interplay between DI hinge restructuring and the N-terminus would rationalize the mechanism of action of many known gB-targeting neutralizing antibodies: the VZV neutralizing antibody 93k^35^ and EBV neutralizing antibody 3A5^34^ may prevent the N-terminus from binding to DIV ^37^, favoring the DI hinge-bound conformation and disrupting the transition to the postfusion conformation.

Conversely, the DI and DII-targeted CMV neutralizing antibodies 1G2 and SM5^32,63^, as well as the N-terminus-directed CMV neutralizing TRL345-like antibodies^60^, may block the N-terminus from binding the DI hinge and promote premature fusogenic triggering. Given that the prefusion-stabilized EBV gB N-terminus was not resolved bound to the DI hinge and that the CMV gB N-terminus in the DIV-bound conformation, virus- or condition-specific conformations and functions for the N-terminus remain to be investigated.

The generalizability of the prefusion-stabilizing mutations identified here to alpha-, beta-, and gamma-Herpesvirus gB trimers indicate that the underlying molecular mechanism of gB mediated fusion is conserved across Herpesviruses. The observed conservation of structural features among other class III fusion proteins is consistent with a shared fusion mechanism, suggesting possible applicability of these mutations or of the design strategy to even more distant pathogens. Identifying a harmonized mechanism across class III fusion proteins positions us more favorably to efficiently develop stabilized class III fusion protein vaccine candidates for pandemic preparedness. Similar to the addition of prefusion-stabilizing proline mutations to the central helix of class I fusion proteins^45,49,78^, extrapolation of the strategies used to stabilize prefusion gB to more distantly related class III fusion proteins may prove invaluable for vaccine development across several viral families.

## Methods

### Computational stabilization of gB and design of oP3h and I5350A2h

Alphafold2^54^ was used to predict the trimeric prefusion conformation structure of EBV gB using the prefusion conformation structure of CMV gB^40^ as a template. Using this structure, ProteinMPNN^53^ and Disulfide by Design 2.0^79^ were used in parallel to identify stabilizing mutations, including the A293P and Q527C-E634C disulfide bond mutations in EBV gB. The relatively soluble HSV1 fusion loop residues were initially used in EBV gB ectodomains as previously described ^80^, but were replaced with ProteinMPNN^53^ optimized fusion loop mutations subsequent to the addition of I5350A2h.

To design an ultrastable trimer that was narrow and short enough to fit partly in the interior of gB and not clash with additional stabilizing trimers, we iteratively cycled the coordinates of the GCN4 trimer (PDB 4dme) between ProteinMPNN and Alphafold2^53,54^ to optimize the design until convergence on a three-helix bundle (designated optimized parallel 3-helix bundle, herein oP3h) harboring the RINEIER heptad repeat motif. These repeat sequences were predicted to tightly pack isoleucine side chains in the coiled-coil core with interdigitating arginine, glutamate, and asparagine side chains forming a network of salt bridges and hydrogen bonds at the periphery. During optimization, the heptad motif was adjusted to RINAIET to increase the surface presentation of hydrophobic moieties, anticipating hydrophobic interactions with the air-water interface may favor cryoEM particle views perpendicular to the C3 symmetry axis. To add oP3h to the coiled coils of SARS-CoV-2 S and EBV gB, the fusion protein coiled coil <u>a</u>bc<u>d</u>efg repeats (where a and d are hydrophobic residues) were first matched to oP3h. While not necessary for all applications of oP3h, a smoother transition between the coiled coil of the fusion protein and oP3h was accomplished using mutations identified in ProteinMPNN^53^ and screened in Alphafold2^54^ before adding to ectodomain constructs.

To redesign the N- and C-termini of I5350A to create I5350A2h, RFDiffusion^81^ was first used to place de novo backbone residues with a regular secondary structure; cycling between ProteinMPNN and Alphafold2^53,54^ was then used to identify residues that would promote formation of this structure. This enabled creating a helix-turn-helix, antiparallel coiled coil, and β-strands that match strands of gB DI to I5350A. Similar to the repeats of oP3h, the use of a coiled coil in this design enabled extension of the structured link between gB and I5350A2h simply by adding heptad repeats: RIQELER repeats to the N-terminus of I5350A2h plus RLREEIN repeats to the C-terminus of I5350A2h. To further redesign I5350A2h to create I5350A2hN, the coordinates of EBV gB-deliDoP-I5350A2h were used with ProteinMPNN to optimize the coiled coil residues. Next, RFDiffusion^81^ was used to place the β-strands that match strands of gB DI in a direction ∼30° tilted from the corresponding residues in I5350A2h. In the context of trimeric I5350A2h these residues point away from the C3 symmetry axis, while they point ∼30° more parallel to this axis in I5350A2hN; consequently I5350A2hN was anticipated to position DI with a narrower diameter than I5350A2h. Cycling between ProteinMPNN and Alphafold2^53,54^ was then used to identify residues that would promote formation of this structure.

### Recombinant ectodomain production

All ectodomains were produced in 25 mL culture of Expi293F Cells (ThermoFisher Scientific) grown in suspension using Expi293 Expression Medium (ThermoFisher Scientific) at 37°C in a humidified 8% CO2 incubator rotating at 130 rpm). Expi293F is a derivative cell line coming from the 293 cell line (Thermo Fisher). Cells grown to a density of 3 million cells per mL were transfected with the ExpiFectamine 293 Transfection Kit (ThermoFisher Scientific) and cultivated for four days at which point the supernatant was harvested. His-tagged ectodomain was purified from clarified supernatants using 2 mL of cobalt resin (Takara Bio TALON) plus 18 µl 1 M CoCl_2_ to compensate for cobalt leaching away with the Expi293 Expression Medium, washing with 200 column volumes of 50 mM Tris-HCl pH 8.0, 150 mM NaCl, and 5 mM imidazole, and eluted with Tris-HCl pH 8.0, 150 mM NaCl, and 600 mM imidazole. Ectodomains were concentrated and buffer exchanged with Tris-HCl pH 8.0 and 150 mM NaCl using a 100 kDa centrifugal filter (Amicon Ultra 0.5 mL centrifugal filters, MilliporeSigma). At this state, ectodomains were evaluated by negative stained EM (see below). Ectodomains selected for cryoEM analysis were further purified by size exclusion chromatography using a Superose 6 increase 10/300 GL column (Cytiva) equilibrated in a buffer containing Tris-HCl pH 8.0 and 150 mM NaCl, and concentrated using a 100 kDa centrifugal filter (Amicon Ultra 0.5 mL centrifugal filters, MilliporeSigma), and flash frozen with liquid nitrogen.

### Negative-stain EM sample preparation

All constructs in this study were negatively stained at a final concentration of 0.01 mg ml^−1^ using Gilder Grids overlaid with a thin layer of carbon and 2% uranyl formate. Data were acquired using the Leginon software^82^ to control a Tecnai T12 transmission electron microscope operated at 120 kV and equipped with a Gatan 4K Ultrascan CCD detector. The dose rate was adjusted to 50 electrons per Å^2^ and each micrograph was acquired in 1 s. For each ectodomain, ∼100 micrographs were collected with a defocus range between −1.0 and −2.5 μm. Data were subsequently processed using CryoSPARC^83^.

### CryoEM sample preparation and data collection

To prepare the CryoEM grids for all ectodomains, the following concentrations of proteins with (or without as indicated) detergent were prepared: 5 mg/mL EBV gB-deli with 0.02 % (w/v) fluorinated octyl-maltoside (FOM, Anatrace) detergent, 2 mg/mL EBV gB-Δ, 2 mg/mL EBV gB-deD, 0.1 mg/mL EBV gB-deliDoP-I5350A, 3 mg/mL EBV gB-deliDoP-I5350A2h with 0.01 % (w/v) FOM, 5 mg/mL EBV gB-deliDoP-I5350A2hN with 0.02 % (w/v) FOM, 3 mg/mL CMV gB-deliDoP-I5350A2h with 0.01 % (w/v) FOM, 3 mg/mL HHV6B gB-deliDoP-I5350A2h with 0.01 % (w/v) FOM, 0.5 mg/mL HSV1 gB-deliD, 1.25 mg/mL VZV gB-deliD + L689K, 1 mg/mL SARS-CoV-2 S-2P + 3X-RINEIER, and 1 mg/mL SARS-CoV-2 S-2P + 4X-RINAIET. FOM was added as indicated to reduce issues of preferred orientation or unfolding at the air water interface. These solutions (3 µL) were added onto a freshly glow discharged 2.0/2.0 UltraFoil^84^ grid (200 mesh), and plunge frozen using a vitrobot MarkIV (ThermoFisher Scientific) using a blot force of −1 and 6 second blot time at 100% humidity and 23°C. For EBV gB-deliDoP-I5350A, prior to glow discharging the 2.0/2.0 UltraFoil^84^ grid (200 mesh) was manually overlaid with a thin layer of carbon. Data were acquired using the Leginon software^82^ to control an FEI Titan Krios transmission electron microscope operated at 300 kV equipped with a Gatan K3 Summit direct detector and Gatan Quantum GIF energy filter, operated in zero-loss mode with a slit width of 20 eV. The dose rate was adjusted to 15 counts/pixel/s, and each movie was acquired in 75 frames of 40 ms with pixel size 0.83 Å. Data were collected with a defocus range between −0.5 and −2.8 μm. For EBV gB-deliDoP-I5350A, HSV1 gB-deliD, SARS-CoV-2 S-2P + 4X-RINEIER, and SARS-CoV-2 S-2P + 4X-RINAIET data were acquired using the Leginon software^82^ to control a Glacios transmission electron microscope equipped with a Gatan K3 Summit direct detector and operated at 200 kV. The dose rate was adjusted to 7.5 counts/pixel/s, and each movie was acquired in 100 frames of 50 ms with pixel size 0.89 Å. For EBV gB-deD, 3141 movies were collected at 0° tilt, 3712 movies were collected at 15° tilt, 3783 movies were collected at 30° tilt, and 519 movies collected at 45° tilt. For HSV gB-deliD and VZV gB-deliD + L689K, data was collected at 30° tilt. All other datasets were collected as a single session without tilt.

### Overall CryoEM data processing

For all datasets, movie frame alignment and pixel binning by 2-fold was initially carried out using Warp^85^, except for EBV gB-deD and EBV gB-deliDoP-I5350A, for which movie frame alignment was performed in CryoSPARC^83^ and 2-fold binning was performed at the particle extraction stage (for these datasets the maps were resolved better this way). For all datasets, estimation of the microscope contrast-transfer function parameters using Patch CTF, particle picking using Topaz^86^, and particle extraction was carried out in CryoSPARC^83^. Reference-free 2D classification was performed using CryoSPARC to select well-defined particle images. Ab initio structure reconstruction and non-uniform refinement in CryoSPARC were then performed, followed by 3D classification in Relion^87^ for EBV gB-deli, SARS-CoV-2 S-2P + 4X-RINEIER, SARS-CoV-2 S-2P + 4X-RINAIET, and VZV gB-deliD + L689K datasets. Alternatively, for EBV gB-deliD, EBV gB-deliDoP-I5350A2h, EBV gB-deliDoP-I5350A2hN, CMV gB-deliDoP-I5350A2h, HHV6B gB-deliDoP-I5350A2h, and HSV1-deliD datasets, ab initio structure reconstruction with multiple classes followed by heterogeneous refinement was performed in CryoSPARC, as this yielded better resolved maps for these datasets. At this stage, particles were subjected to reference-based motion correction in CryoSPARC^83^ (for datasets with movie frames aligned in CryoSPARC^83^) or Bayesian polishing using Relion^88^ (for datasets with movie frames aligned in WARP) during which the pixels were unbinned. Reference-based motion correction/Polishing was not performed on the EBV gB-Δ and EBV gB-deliDoP-I5350A datasets, as this did not yield a better resolved maps. Another round of non-uniform refinement in CryoSPARC was performed concomitantly with global and per-particle defocus refinements as well as beam tilt refinement, parameters which were optimized for each dataset^87,89^. Reported resolutions are based on the gold-standard Fourier shell correlation (FSC) of 0.143 criterion and Fourier shell correlation curves were corrected for the effects of soft masking by high-resolution noise substitution^90,91^.

### CryoEM data processing for local features

For SARS-CoV-2 S-2P + 4X-RINEIER and SARS-CoV-2 S-2P + 4X-RINAIET datasets, a mask surrounding oP3h was generated in UCSF Chimera^92^, and used for local refinement of this region applying C3 symmetry in CryoSPARC^83^. For HSV1 gB deliD, map resolvability was best when all-open particles and all-closed particles were separated, but not when trimers and dimers of trimers were separated. UCSF Chimera^92^ was used to create a mask surrounding the region corresponding to an individual trimer, which was used for local refinement of this region applying C3 symmetry in CryoSPARC^83^.

For refining the map of I5350A2h from EBV gB-deliDoP-I5350A2h, masks were created in UCSF Chimera^92^ surrounding the gB ectodomain and I5350A2h, these masks were used for multibody refinement followed by particle signal subtraction in Relion^93^ to remove signal from the gB ectodomain, and then the mask surrounding I5350A2h was used for local refinement of this region applying C3 symmetry in CryoSPARC^83^. For refining the map of DII-DV from EBV gB-deliDoP-I5350A2h, masks were created in UCSF Chimera^92^ surrounding the DII-DV from the gB ectodomain and DI with I5350A2h, these masks were used for multibody refinement followed by particle signal subtraction in Relion^93^ to remove signal from DI and I5350A2h, and then the mask surrounding DII-DV was used for local refinement of this region in CryoSPARC^83^. For refining the map of DIII-DV from EBV gB-deliDoP-I5350A2h, a mask of this region was created in UCSF Chimera^92^ and used to locally refine the particles from the local refinement of DII-DV above, applying C3 symmetry in CryoSPARC^83^. DI is framed by two semi-flexible pivot points and remains poorly resolved in EBV gB-deliDoP-I5350A2h maps, and it is too small for local refinement strategies on its own.

For locally refining the map of EBV gB-deliDoP-I5350A2hN, masks were created in UCSF Chimera^92^ surrounding the gB ectodomain and I5350A2hN, these masks were used for multibody refinement followed by particle signal subtraction in Relion^93^ to remove signal from the gB ectodomain or I5350A2hN. Then a mask surrounding I5350A2hN or gB, was used for local refinement of this region applying C3 or C1 symmetry, respectively, in CryoSPARC^83^, yielding maps for I5350A2hN and the gB ectodomain. Finally, to improve the local resolution of DII-DV of EBV gB-deliDoP-I5350A2hN, a mask of this region was created in UCSF Chimera^92^ and used to locally refine the particles from the local refinement of DI-DV above, applying C3 symmetry in CryoSPARC^83^.

For refining the map of DII-DV from CMV gB-deliDoP-I5350A2h, masks were created in UCSF Chimera^92^ surrounding the DII-DV from the gB ectodomain and DI with I5350A2h, these masks were used for multibody refinement followed by particle signal subtraction in Relion^93^ to remove signal from DI and I5350A2h, and then the mask surrounding DII-DV was used for local refinement of this region in CryoSPARC applying C3 symmetry^83^. As for EBV above, to modestly improve resolution of DI, UCSF Chimera^92^ was used to create a mask surrounding DII-DV plus DI from two protomers and a mask surrounding the last DI with I5350A2h; these masks were used for multibody refinement followed by particle signal subtraction in Relion^93^ to remove signal from I5350A2h, and then the mask surrounding DII-DV plus DI from two protomers was used for local refinement of this region in CryoSPARC^83^.

For refining the map of DII-DV from HHV6B gB-deliDoP-I5350A2h, masks were created in UCSF Chimera^92^ surrounding the DII-DV from the gB ectodomain and DI with I5350A2h, these masks were used for multibody refinement followed by particle signal subtraction in Relion ^93^ to remove signal from DI and I5350A2h, and then the mask surrounding DII-DV was used for local refinement of this region in CryoSPARC applying C3 symmetry^83^. Though the ‘crown’ portion of DIII, DIV, and DV is visible in low-pass filtered maps, this region was not resolved otherwise, even in local refinements without applying symmetry.

### Cryo-EM model building and analysis

For VZV gB-deliD, EBV gB-deli, EBV gB-deD, EBV gB-deliDoP-I5350A2h, and EBV gB-deliDoP-I5350A2hN maps, EMready^94^ was used to improve map quality for model building; otherwise the unsharpened maps and locally sharpened maps were used for model refinement and deposition. Initial models of ectodomains were predicted using Alphafold2^54^ and rigidly docked into corresponding maps in UCSF Chimera^92^. Models were then refined and rebuilt by iteratively cycling between Coot, ISOLDE, and Rosetta: Briefly, self-restraints between atoms in these models were created to maintain structured features prior to flexibly fitting the model coordinates to the map in Coot^95,96^. At this stage, ISOLDE^97^ (implemented in ChimeraX^98^) was used to relax the model into the map with an AMBER forcefield and remodel as necessary. Rosetta^99,100^ was then used to further optimize fit and geometry. Models were validated using MolProbity^101^, EMringer^102^, and Phenix^103^. Figures were generated using UCSF ChimeraX^98^ and UCSF Chimera^92^.

---

**Table S2.**
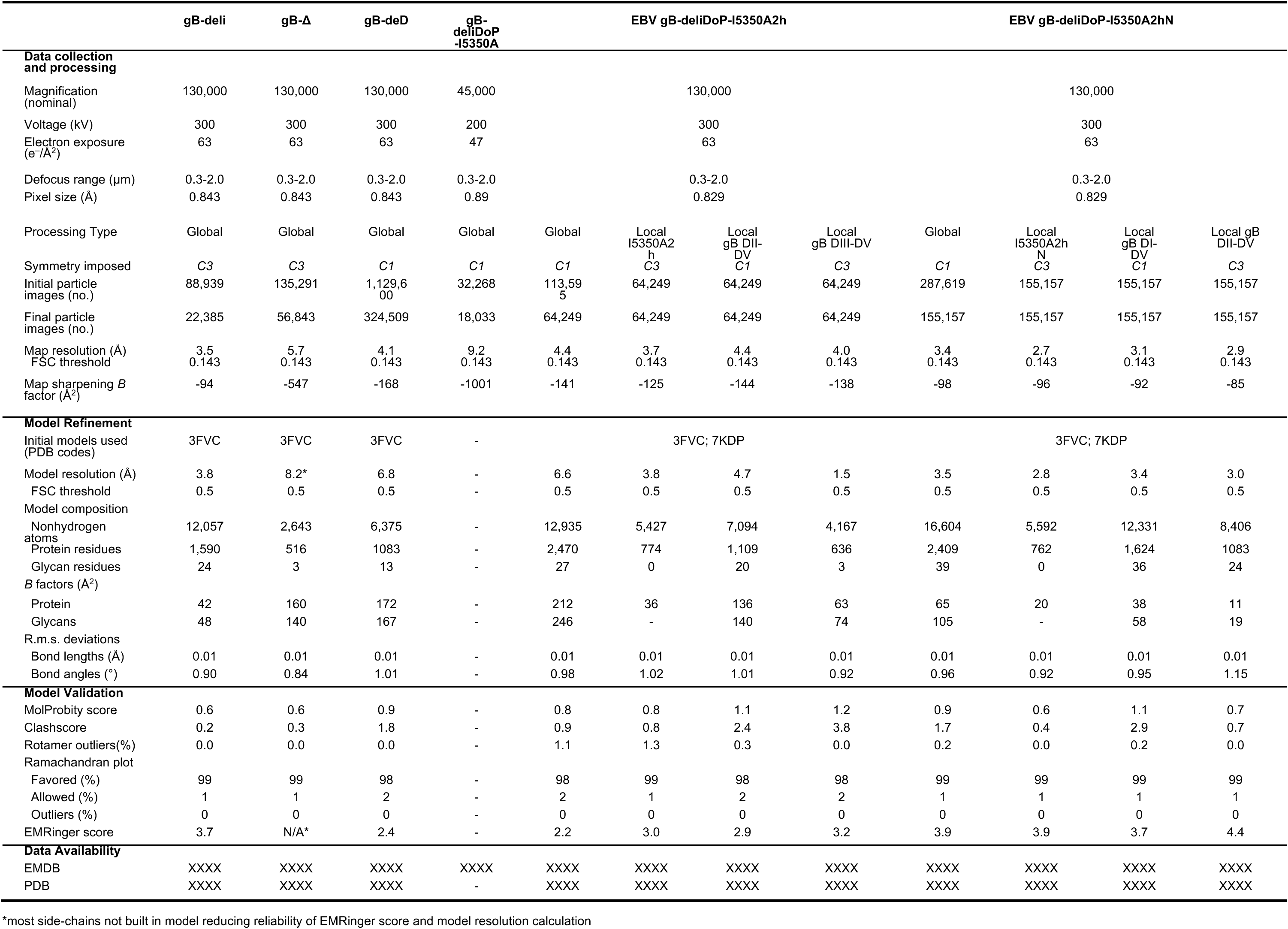
Cryo-EM data collection, refinement and validation statistics for EBV gB constructs.

**Table S3.**
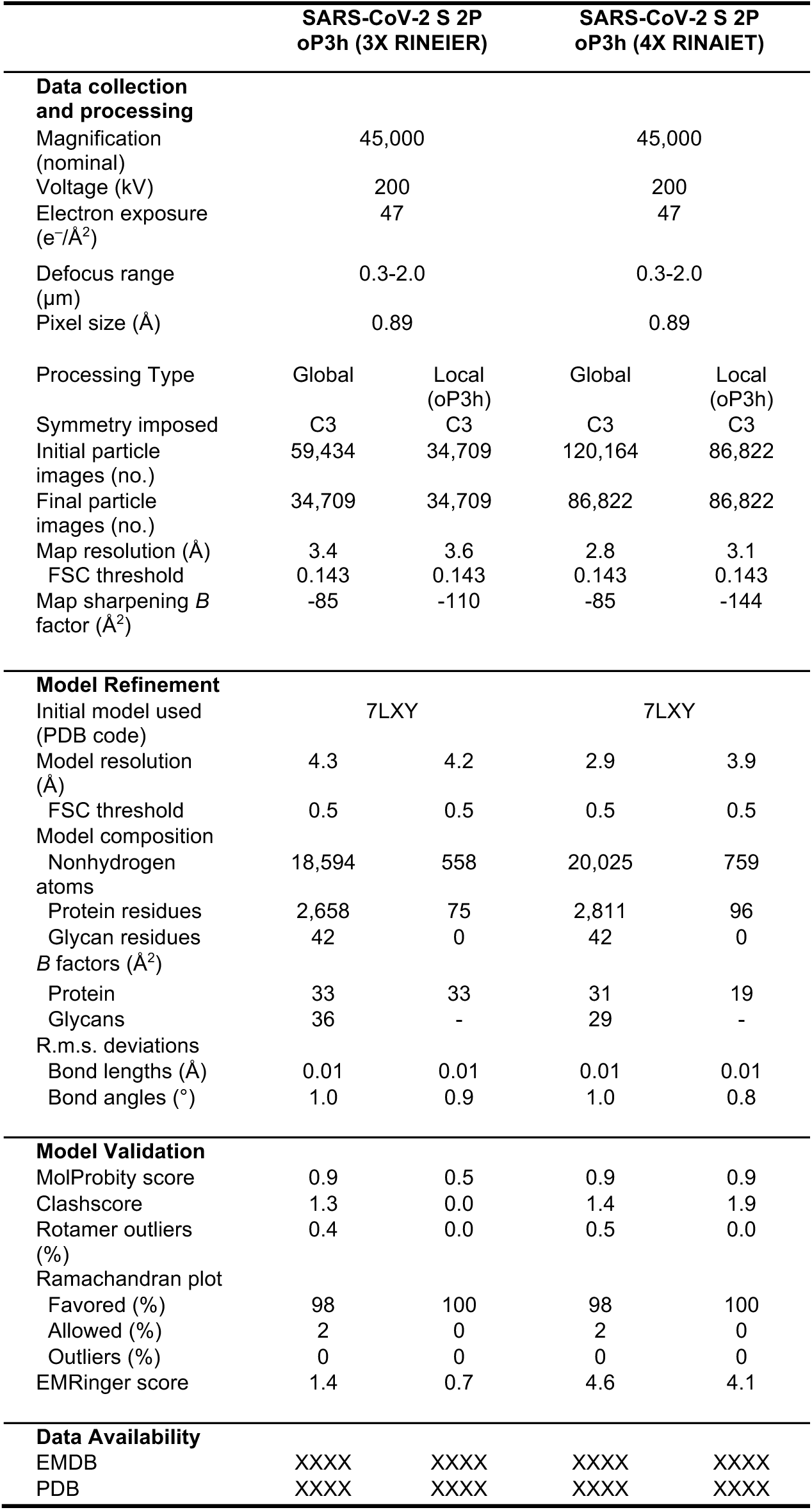
Cryo-EM data collection, refinement and validation statistics for SARS-CoV-2 S 2P oP3h constructs.

**Table S4.** Cryo-EM data collection, refinement and validation statistics for CMV, HHV6B, HSV1, and VZV gB constructs.

|  | CMV gB-deliDoP<br>I5350A2a |  | HHV6B gB-deliDoP<br>I5350A2a |  | HSV1 gB-deliD |  | VZV gB-deliD |
| --- | --- | --- | --- | --- | --- | --- | --- |
| Data collection and processing |  |  |  |  |  |  |  |
| Magnification (nominal) | 130,000 |  | 130,000 |  | 45,000 |  | 130,000 |
| Voltage (kV) | 300 |  | 300 |  | 200 |  | 300 |
| Electron exposure (e<sup>-</sup>/Å<sup>2</sup>) | 63 |  | 63 |  | 47 |  | 63 |
| Defocus range (μm) | 0.3-2.0 |  | 0.3-2.0 |  | 0.3-2.0 |  | 0.3-2.0 |
| Pixel size (Å) | 0.829 |  | 0.829 |  | 0.89 |  | 0.843 |
| Processing Type | Global | Local<br>gB DII-DV | Global | Global | Local<br>(closed) | Local<br>(open) | Global |
| Symmetry imposed | C1 | C3 | C3 | C3 | C3 | C3 | C3 |
| Initial particle images (no.) | 51,511 | 33,533 | 85,490 | 67,753 | 138,213 | 138,213 | 251,305 |
| Final particle images (no.) | 33,533 | 33,533 | 67,753 | 67,753 | 43,635 | 40,021 | 48,271 |
| Map resolution (Å) | 6.2 | 5.8 | 4.3 | 4.6 | 5.8 | 6.3 | 2.9 |
| FSC threshold | 0.143 | 0.143 | 0.143 | 0.143 | 0.143 | 0.143 | 0.143 |
| Map sharpening *B* factor (Å<sup>2</sup>) | -461 | -455 | -113 | -251 | -362 | -442 | -96 |
| Model Refinement |  |  |  |  |  |  |  |
| Initial model used (PDB code) | 7KDP |  | 7KDP |  | 6Z9M |  | 7K1S |
| Model resolution (Å) | 7.9* | 7.2* | 4.5* | 6.7* | 8.4* | 8.4* | 3.3 |
| FSC threshold | 0.5 | 0.5 | 0.5 | 0.5 | 0.5 | 0.5 | 0.5 |
| Model composition |  |  |  |  |  |  |  |
| Nonhydrogen atoms | 13,049 | 5,802 | 10,055 | 2,747 | 8,451 | 8,538 | 25,656 |
| Protein residues | 2,482 | 1086 | 1920 | 507 | 1650 | 1,632 | 3,342 |
| Glycan residues | 33 | 21 | 25 | 12 | 6 | 18 | 18 |
| *B* factors (Å<sup>2</sup>) |  |  |  |  |  |  |  |
| Protein | 200 | 122 | 300 | 101 | 197 | 196 | 22 |
| Glycans | 233 | 117 | 388 | 100 | 202 | 217 | 25 |
| R.m.s. deviations |  |  |  |  |  |  |  |
| Bond lengths (Å) | 0.01 | 0.01 | 0.01 | 0.01 | 0.01 | 0.01 | 0.01 |
| Bond angles (°) | 1.02 | 0.96 | 1.09 | 1.02 | 0.98 | 1.09 | 0.95 |
| Model Validation |  |  |  |  |  |  |  |
| MolProbity score | 0.6 | 0.8 | 0.8 | 0.7 | 1.3 | 1.2 | 0.9 |
| Clashscore | 0.4 | 0.7 | 1.1 | 0.7 | 1.3 | 1.7 | 1.6 |
| Rotamer outliers (%) | 0.0 | 0.0 | 0.0 | 0.0 | 2.6 | 2.4 | 0.5 |
| Ramachandran plot |  |  |  |  |  |  |  |
| Favored (%) | 98 | 98 | 98 | 99 | 99 | 99 | 99 |
| Allowed (%) | 2 | 2 | 2 | 1 | 1 | 1 | 1 |
| Outliers (%) | 0 | 0 | 0 | 0 | 0 | 0 | 0 |
| EMRinger score | N/A* | N/A* | N/A* | N/A* | N/A* | N/A* | 4.6 |
| Data Availability |  |  |  |  |  |  |  |
| EMDB | XXXX | XXXX | XXXX | XXXX | XXXX | XXXX | XXXX |
| PDB | XXXX | XXXX | XXXX | XXXX | XXXX | XXXX | XXXX |

\*most side-chains not built in model reducing reliability of EMRinger score and model resolution calculation

## Extended Data Figures

**Extended Data Figure 1.**
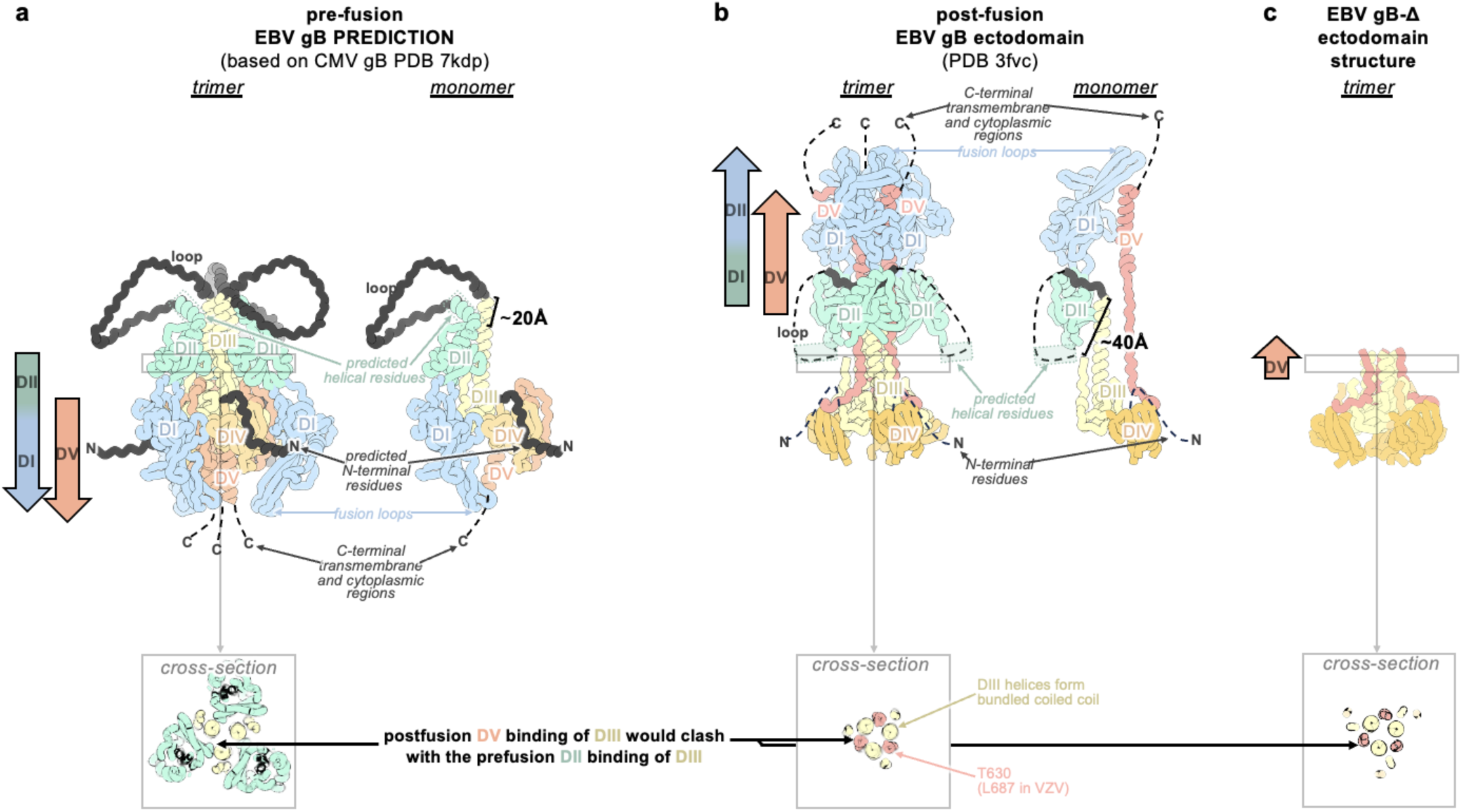
DI, DII, and DV changes between the Alphafold2 predicted prefusion gB (a) postfusion gB (b) and gB-Δ conformations, aligned to the central helices of DIII (c). Panels show the gB trimer (left) or a single protomer (right). Domain I (DI, blue), domain II (DII, green), domain III (DIII, yellow), domain IV (DIV, orange), domain V (DV, red). In the prefusion structure prediction, the loop between DII and DIII as well as the N-terminus (black) are shown; otherwise unmodeled regions are shown as dotted black lines. The distance the loop spans between rigidly structured elements is ∼20 Å in the prefusion conformation and ∼40 Å in the postfusion conformation. Insets show a cross-section through DIII, highlighting the relative positions of DII and DV bound to DIII in the prefusion and postfusion conformations. In the postfusion conformation, DV binds the bundled or closed central helices of DIII and occupies the former prefusion binding site of DII. Residue T630 in EBV gB (L687 in VZV gB) is labelled.

**Extended Data Figure 2.**
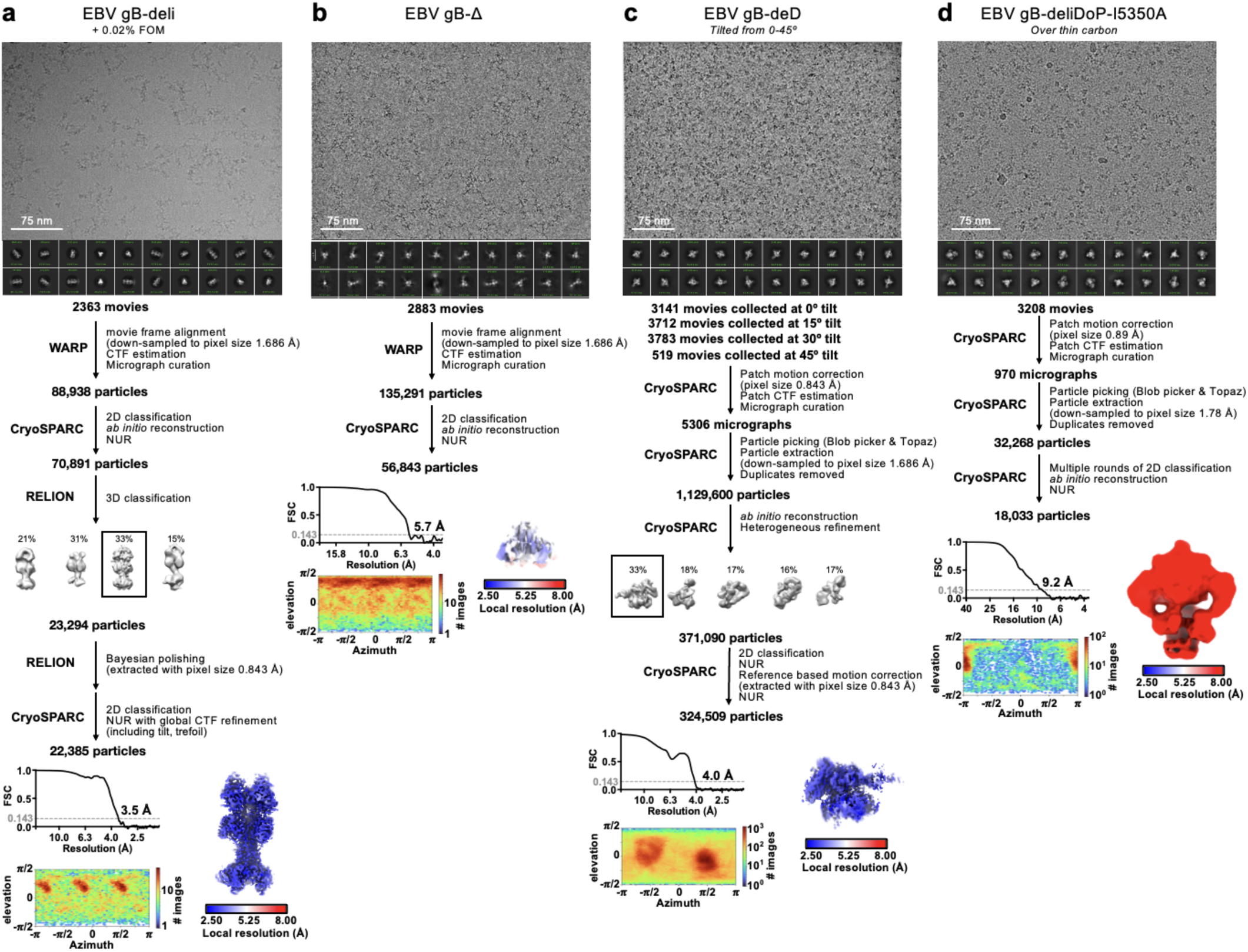
CryoEM data processing flowcharts of EBV gB-deli (a), gB-Δ (b), gB-deliD (c), and gB-deliDoP-I5350A (d) datasets. Representative electron micrograph and 2D class averages are shown at the start of the flowchart. At the end of the flowchart is the gold-standard Fourier shell correlation curve (0.143 cutoff: horizontal dashed line), angular distribution of particles (heat map), and local resolution estimation plotted on the maps (calculated in CryoSPARC). CTF: contrast transfer function; NUR: non-uniform refinement.

**Extended Data Figure 3.**
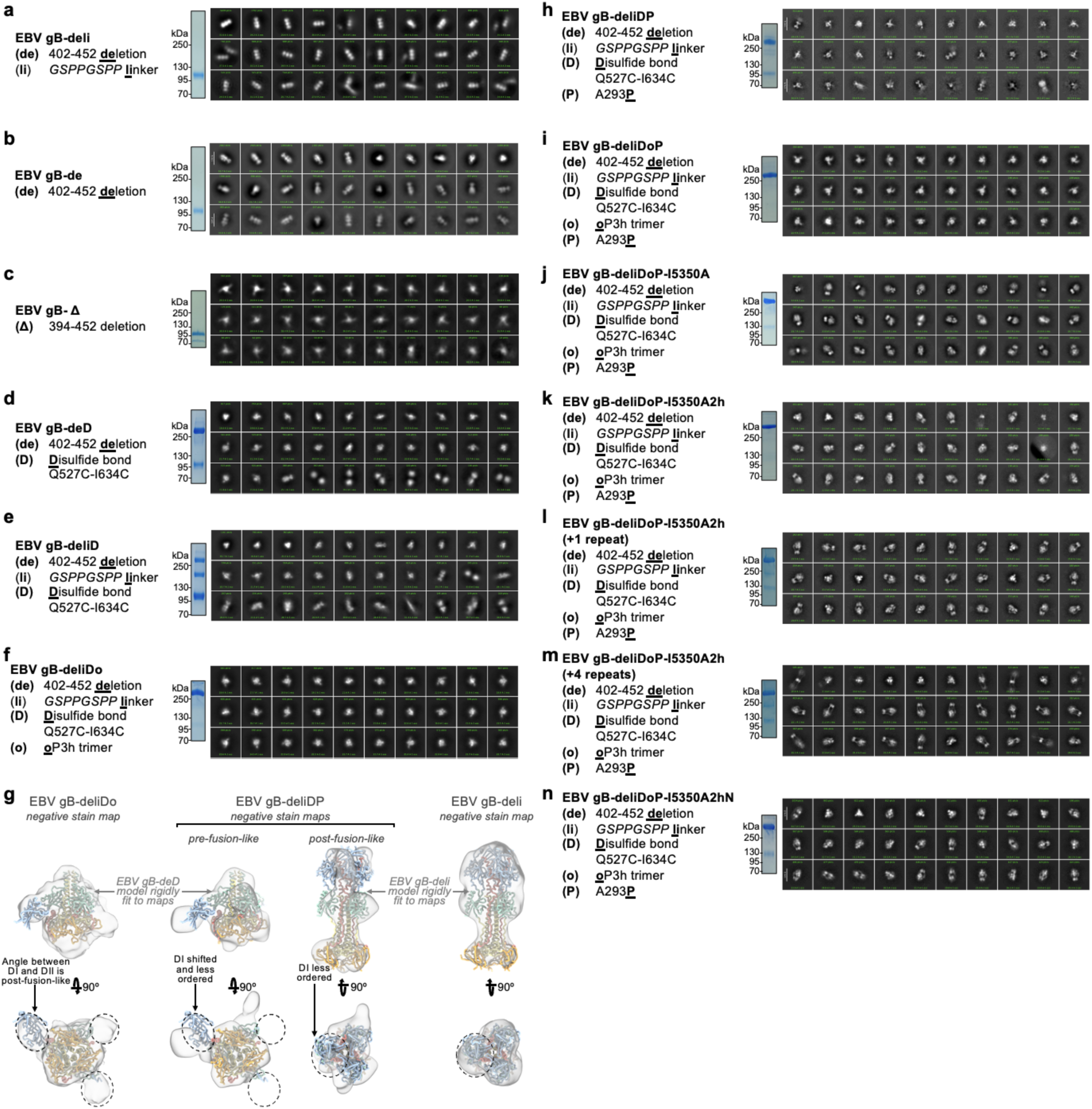
Non-reducing SDS-PAGE, negative stain 2D classes and selected 3D maps of negatively stained EBV gB ectodomain constructs. **(a-f, h-n)** A summary of the construct is listed (left), as well as non-reducing SDS-PAGE (middle), and negative stain 2D classes (right). (**g**) Left, EBV gB-deliDo negative stain map reconstruction with the cryoEM structure of EBV gB-deD (black cartoon) rigidly fit into this map. Center, EBV gB-deliDP negative stain map reconstruction. Right, EBV gB-deli negative stain map reconstruction. Dashed lines highlight the DI orientation in gB-deD, gB-deliDo, and gB-deli, and the apparent movement of DI in EBV gB-deliDP.

**Extended Data Figure 4.**
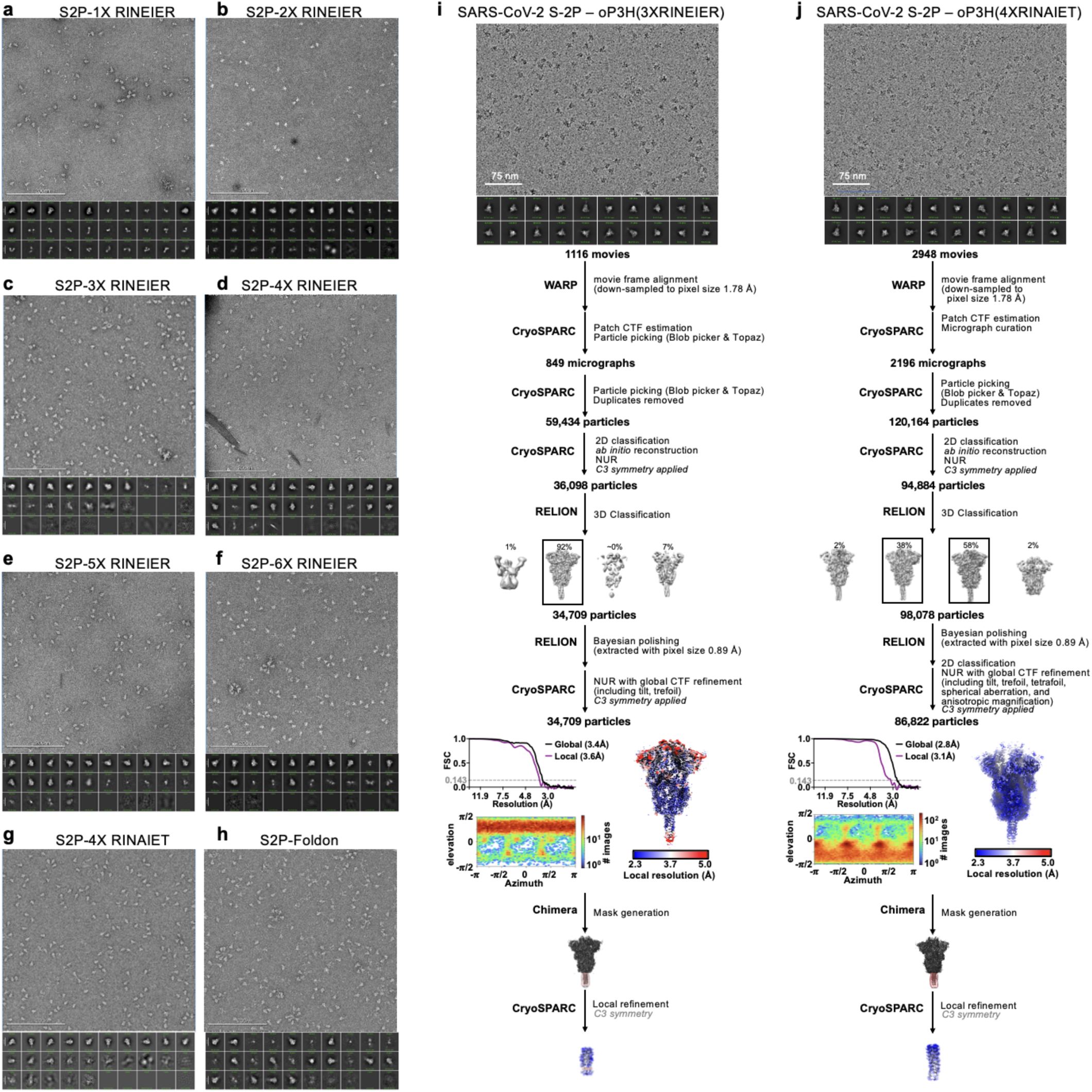
Summary of negative stain EM and cryoEM data for SARS-CoV-2 S-2P – oP3h. **(a-h)** negative stain micrographs (top) and 2D classifications (bottom) of SARS-CoV-2 S-2P with the indicated C-terminal trimerization domains. (**i-j**) CryoEM data processing flowcharts of SARS-CoV-2 S-2P – oP3h (3X RINEIER) (i) and SARS-CoV-2 S-2P – oP3h (4X RINAIET) (j) datasets. At the start of the flowchart, representative electron micrograph and 2D class averages of SARS-CoV-2 S ectodomains embedded in vitreous ice are shown. At the end of the flowchart is the gold-standard Fourier shell correlation curve (0.143 cutoff: horizontal dashed line), angular distribution of particles (heat map), and local resolution estimation plotted on the maps (calculated in CryoSPARC). At the bottom, the local refinement strategies employed for improving map resolution are also shown. CTF: contrast transfer function; NUR: non-uniform refinement.

**Extended Data Figure 5.**
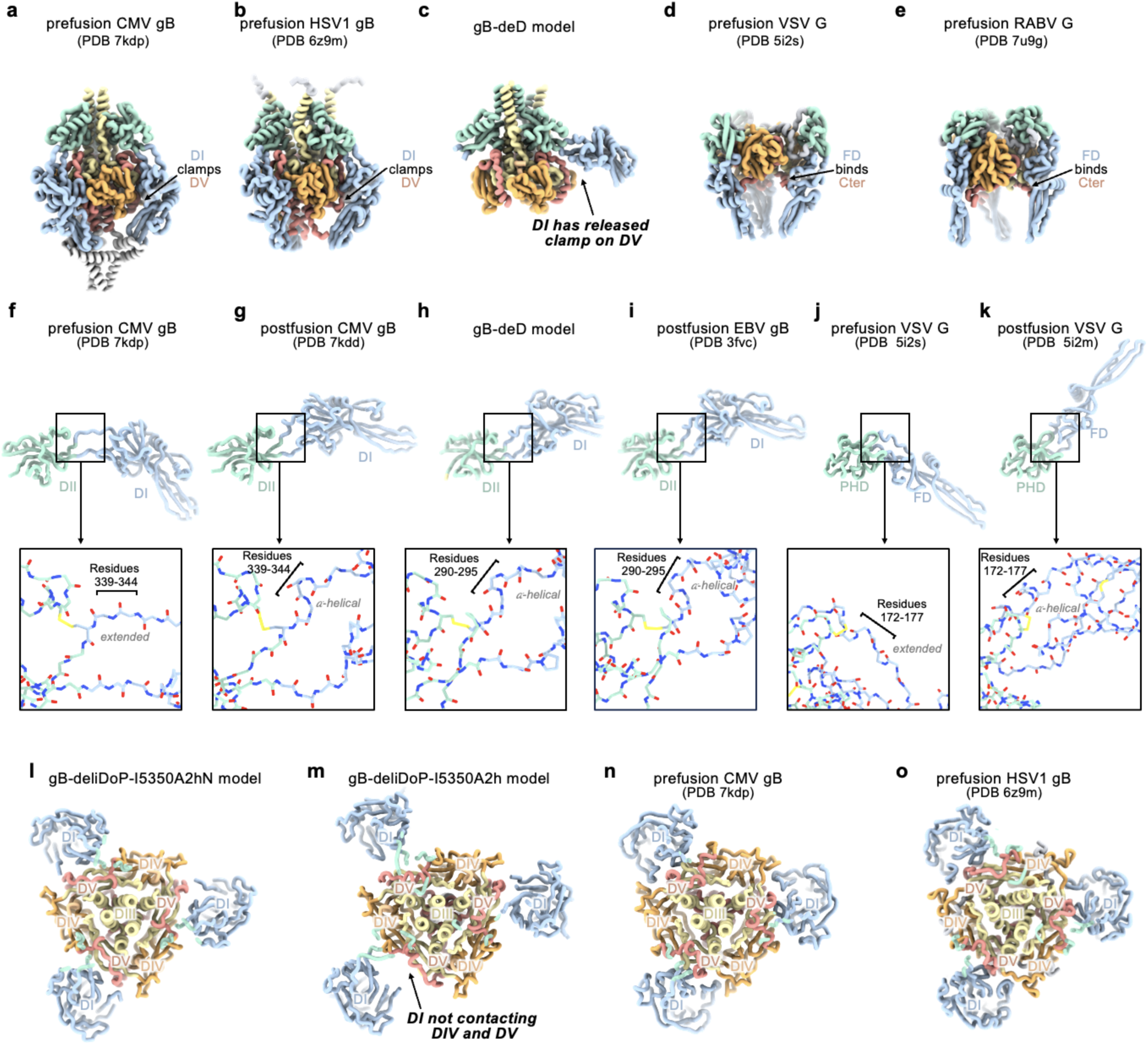
Comparing prefusion and postfusion hinge conformations with gB-deD. Domain I (DI, blue), domain II (DII, green), domain III (DIII, yellow), domain IV (DIV, orange), and domain V (DV, red) are shown as ribbons. (**a-e**) Side-view comparisons of prefusion CMV gB (**a**), prefusion HSV1 gB (**b**), the EBV gB-deD model determined herein (**c**), prefusion VSV G (**d**), and prefusion RABV G (**e**) highlighting how prefusion structures clamp or bind DV (Cter in VSV and RABV G). **(f-k)** Comparisons of DI hinge structure showing DI and DII (or FD and PHD for VSV) from prefusion CMV gB (**f**), prefusion HSV1 gB (**g**), the gB-deD model determined herein (**h**), postfusion EBV gB (i), prefusion VSV G (**j**), and postfusion VSV G (**k**). Insets show zoomed-in view of backbone atoms, plus disulfide bonds and prolines, highlighting the restructuring of prefusion and postfusion DI hinge residues. (**l-o**) Comparisons of gB cross-sections showing DI in relation to DIV and DV, including EBV gB-deliDoP-I5350A2hN (**l**), EBV gB-deliDoP-I5350A2h (**m**), prefusion CMV gB (**n**), and prefusion HSV1 gB (**o**).

**Extended Data Figure 6.**
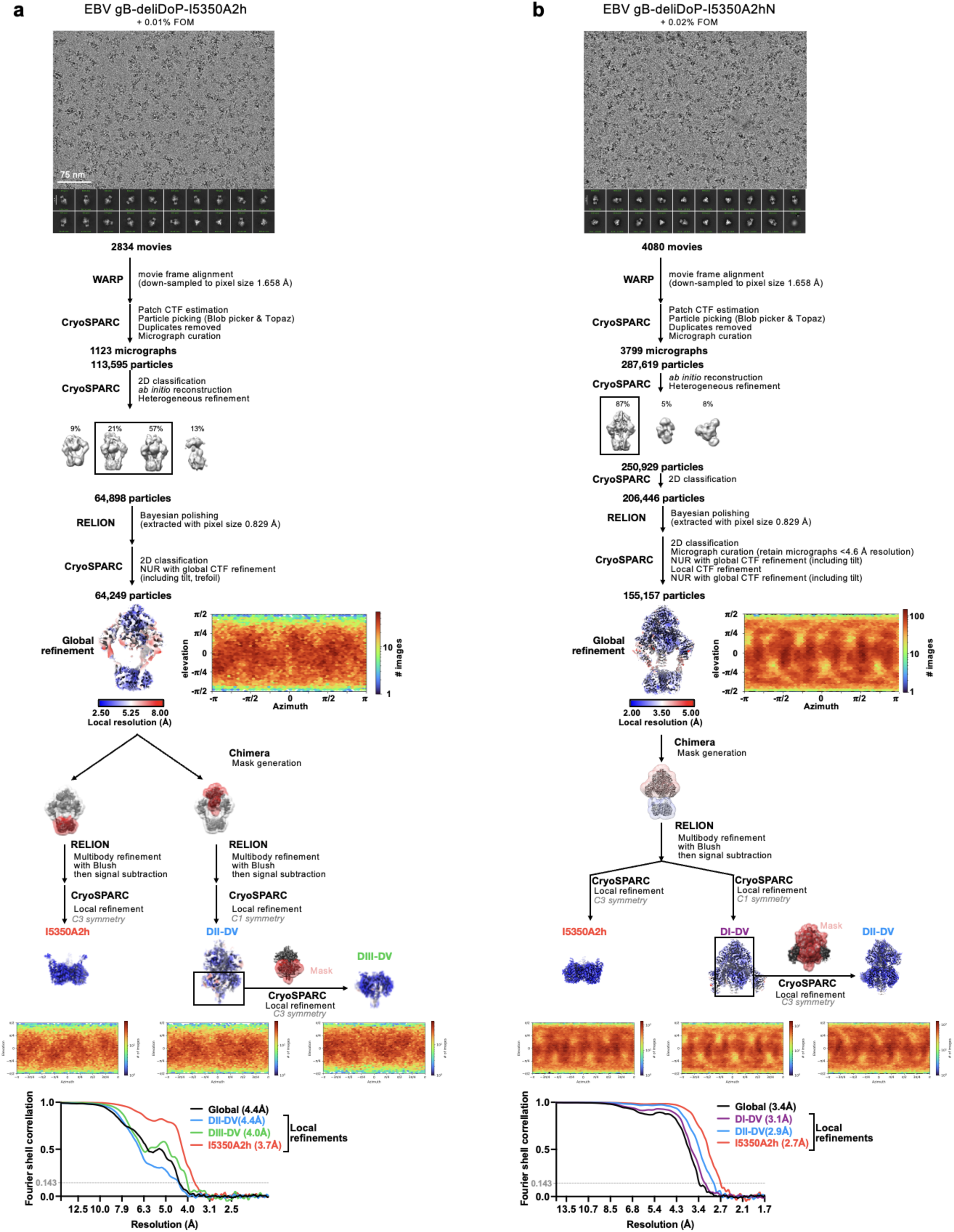
CryoEM data processing flowcharts of EBV gB-deliDoP-I5350A2h (a) and EBV gB-deliDoP-I5350A2hN (b) datasets. Representative electron micrograph and 2D class averages are shown at the start of the flowchart. At the end of the flowchart is the gold-standard Fourier shell correlation curve (0.143 cutoff: horizontal dashed line), angular distribution of particles (heat map), and local resolution estimation plotted on the maps (calculated in CryoSPARC). This flowchart additionally has a summary of local refinement strategies employed for improving local map resolution. CTF: contrast transfer function; NUR: non-uniform refinement.

**Extended Data Figure 7.**
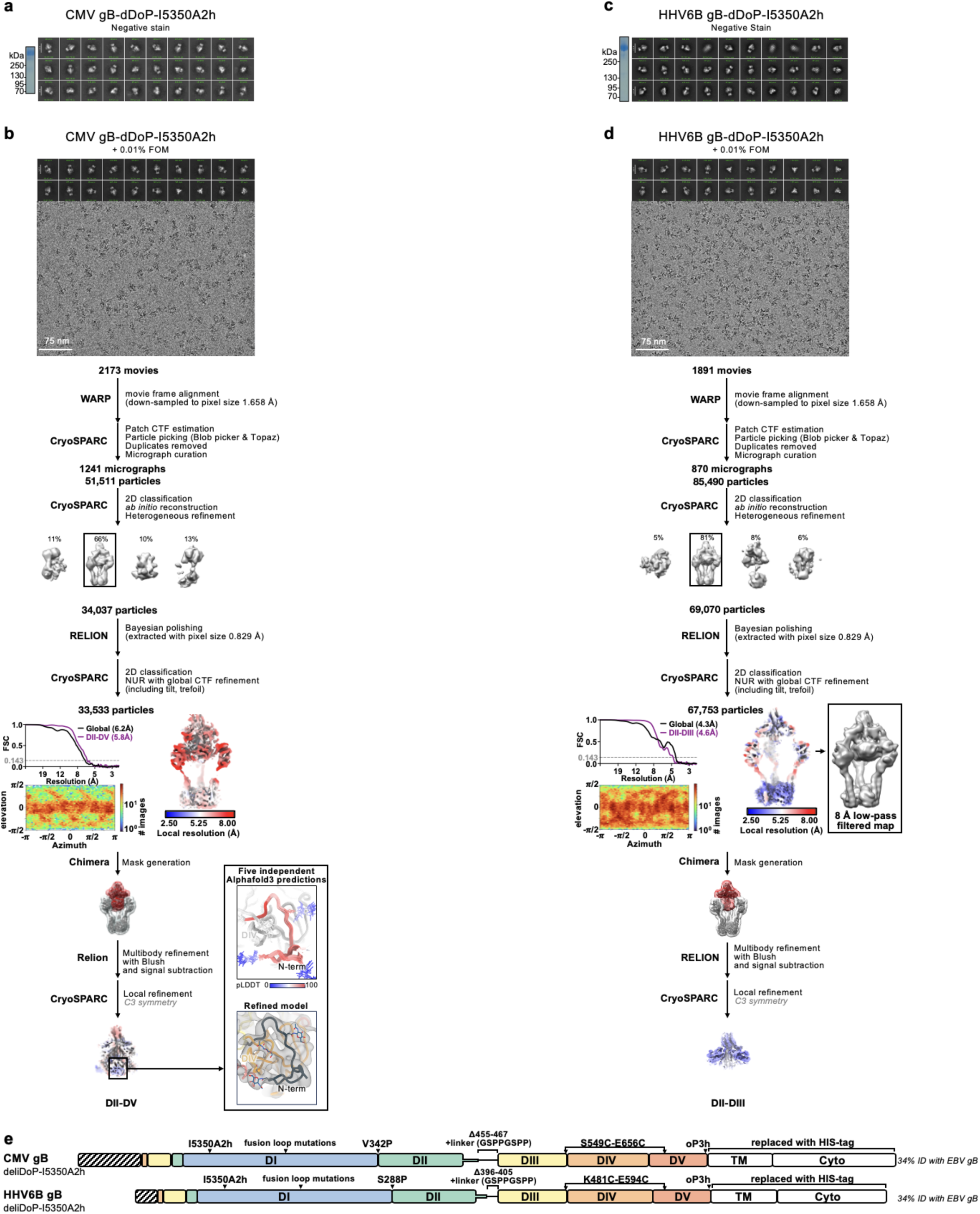
Summary of negative stain EM and cryoEM data for CMV gB-deliDoP-I5350A2h (a-b) and HHV6B gB-deliDoP-I5350A2h (c-d). **(a,c)** non-reducing SDS-PAGE (left) and negative stain 2D classifications (right) of gB-deliDoP-I5350A2h ectodomains. (**c,d**) CryoEM data processing flowcharts of CMV gB-deliDoP-I5350A2h (b), and HHV6B gB-deliDoP-I5350A2h (d) datasets. At the start of the flowchart, representative electron micrograph and 2D class averages of gB ectodomain particles embedded in vitreous ice are shown. At the end of the flowchart is the gold-standard Fourier shell correlation curve (0.143 cutoff: horizontal dashed line), angular distribution of particles (heat map), and local resolution estimation plotted on the maps (calculated in CryoSPARC). At the bottom, the local refinement strategies employed for improving map resolution are also shown. The Alphafold3 predicted structure of the N-terminus of CMV is shown beside the refined model for comparison ^104^. CTF: contrast transfer function; NUR: non-uniform refinement. (**e**) schematic of mutations used for CMV gB-deliDoP-I5350A2h and HHV6B gB-deliDoP-I5350A2h.

**Extended Data Figure 8.**
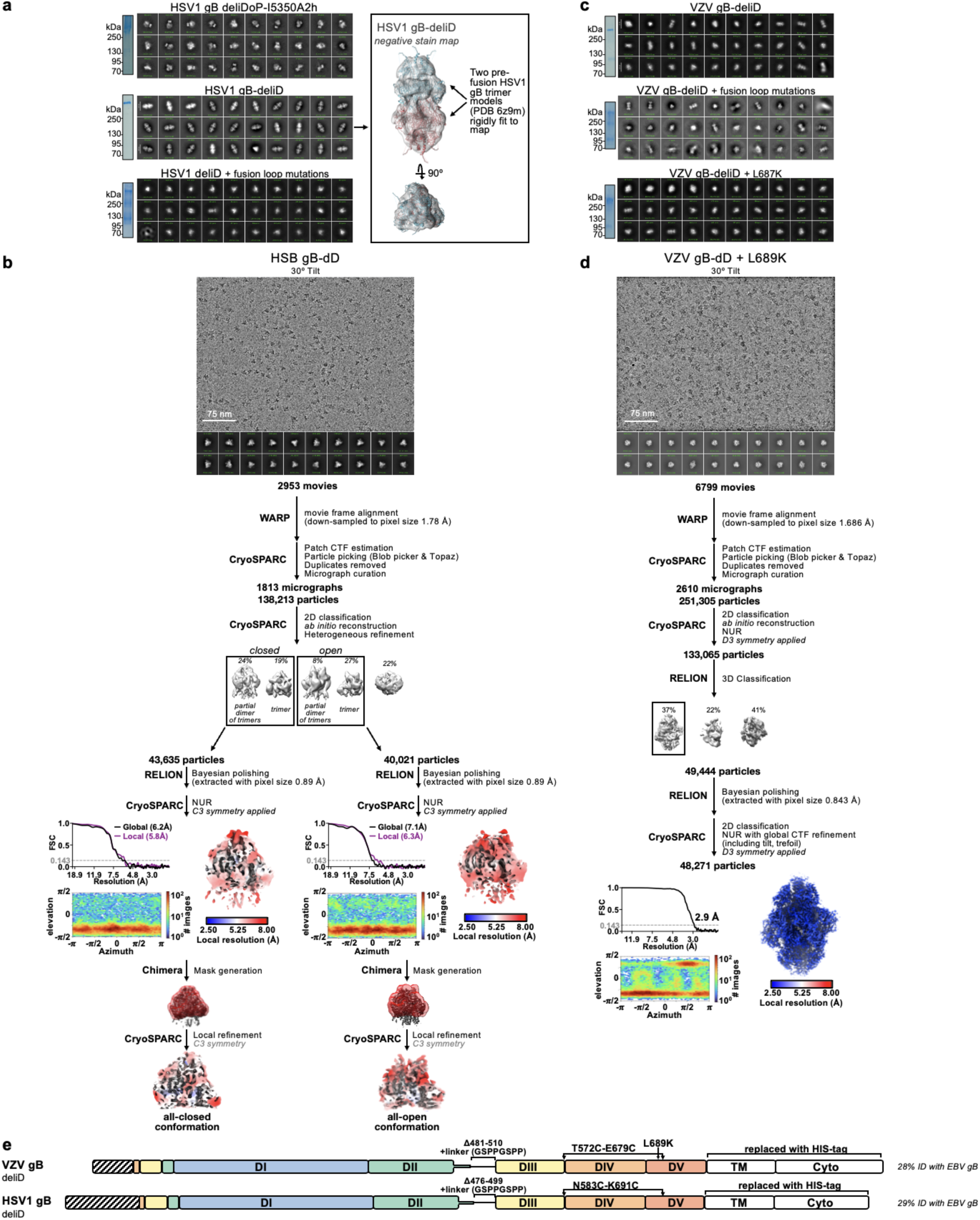
Summary of negative stain EM and cryoEM data for HSV1 and VZV gB ectodomains. **(a,c)** non-reducing SDS-PAGE (left) and negative stain 2D classifications (right) of HSV1 and VZV gB ectodomains. Inset shows HSV1 gB-deliD negative stain map reconstruction with two copies of prefusion HSV1 gB (PDB 6z9m, blue and red ribbons) rigidly fit into the map. (**c,d**) CryoEM data processing flowcharts of HSV1 gB-deliD (**c**), and VZV-deliD + L689K (**d**) datasets. At the start of the flowchart, representative electron micrograph and 2D class averages of gB ectodomain particles embedded in vitreous ice are shown. At the end of the flowchart is the gold-standard Fourier shell correlation curve (0.143 cutoff: horizontal dashed line), angular distribution of particles (heat map), and local resolution estimation plotted on the maps (calculated in CryoSPARC). At the bottom, the local refinement strategies employed for improving map resolution are also shown for HSV-1 gB-deliD. CTF: contrast transfer function; NUR: non-uniform refinement. (**e**) schematic of mutations used for VZV gB-deliD + L689K and HSV1 gB-deliD.

**Extended Data Figure 9.**
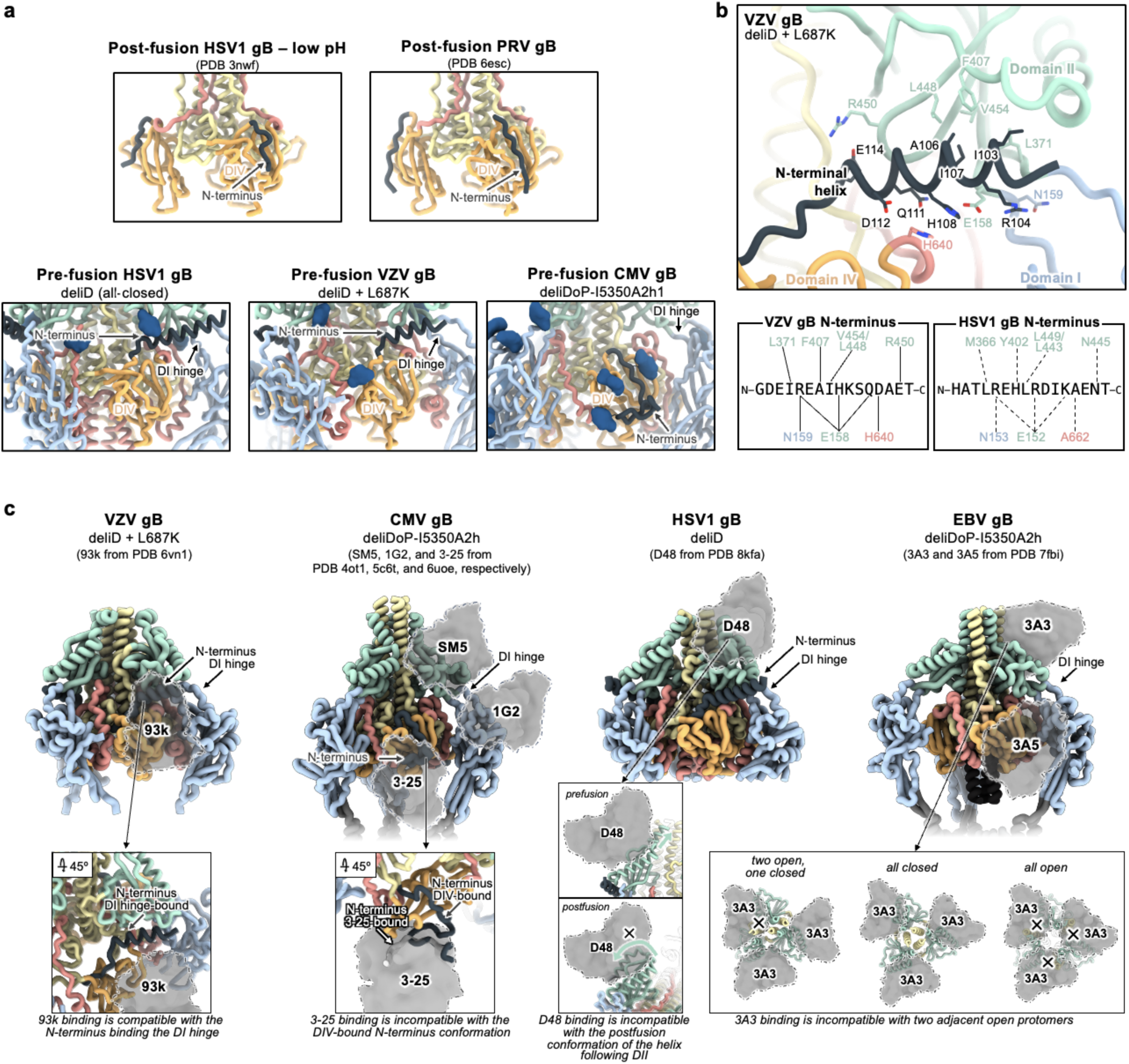
Comparing structurally characterized neutralizing antibody binding sites to prefusion-stabilized gB. (**a**) Structures of gB with the N-terminus (black ribbon) bound to DIV or the DI hinge; Domain I (DI, blue), domain II (DII, green), domain III (DIII, yellow), domain IV (DIV, orange), and domain V (DV, red) are shown as a ribbons. Shown are HSV1 gB, PRV (pseudorabies) gB, HSV1 gB-deliD, VZV gB-deliD, and CMV gB-deliDoP-I5350A2h. (**b**) Zoomed-in view of VZV gB deliD + L687K cryoEM structure, focusing on the N-terminal helix and DI hinge. Domains are colored as in **a** with key residues labeled. The lower left inset shows a schematic of the VZV gB N-terminal residues, with interacting residues. The lower right inset shows a schematic of the HSV1 gB N-terminal residues and residues aligning to the residues shown in the left inset for VZV gB. (**c**) Anti-gB neutralizing antibody structures, with the domains bound by these antibodies aligned to the prefusion conformation determined here, to show the epitopes bound in the prefusion conformation. Antibody variable domains shown as semi-transparent grey surfaces. The lower panels show a zoomed-in view of antibody interfaces, highlighting how these antibodies would constrain structural changes or prevent conformations of gB.

**Extended Data Figure 10.**
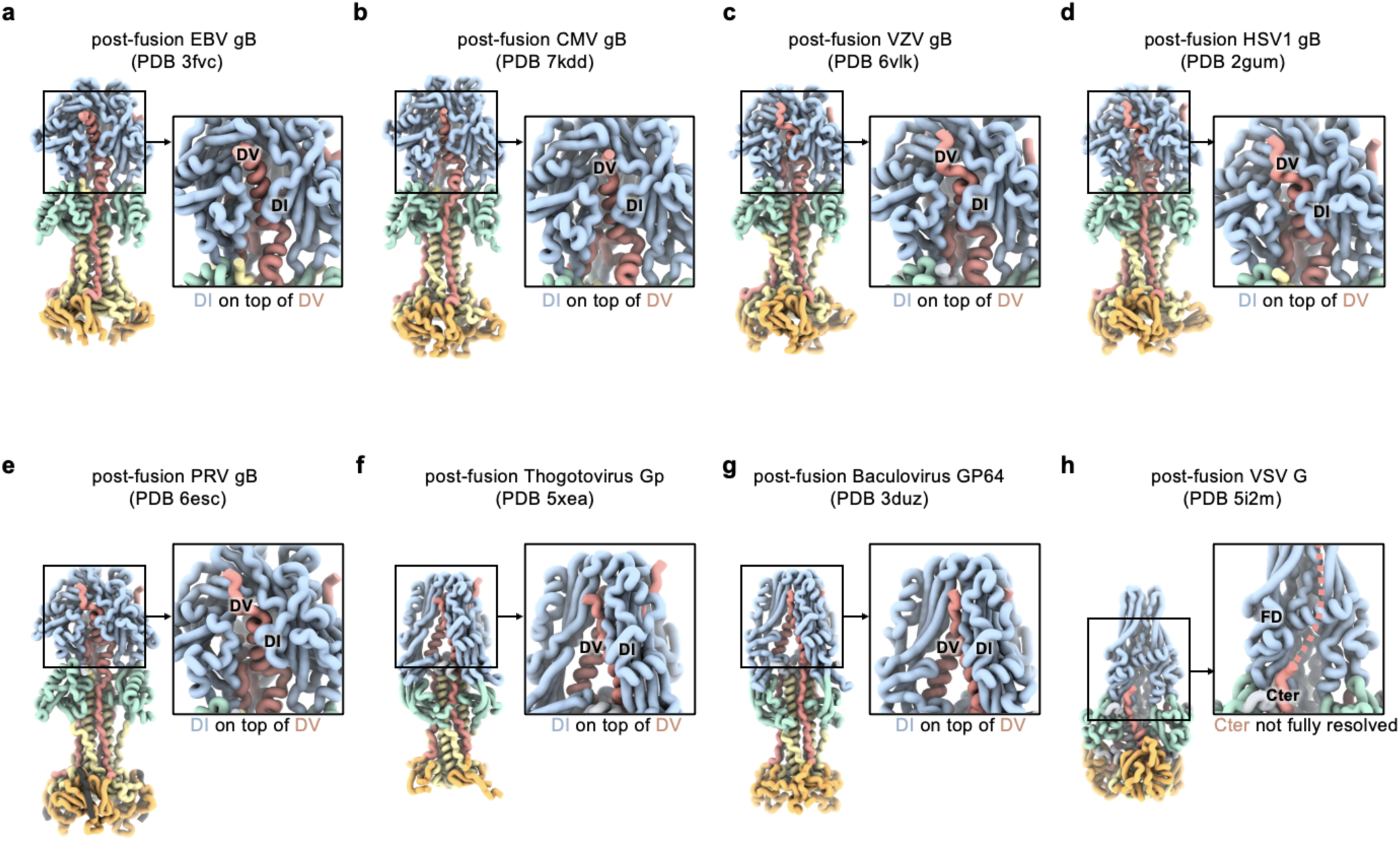
Comparison of class III fusion protein postfusion structures. Postfusion structures shown as ribbons for EBV gB (**a**), CMV gB (**b**), VZV gB (**c**), HSV1 gB (**d**), PRV gB (**e**), Thogotovirus Gp (**f**), Baculovirus GP64 (**g**), and VSV G (**h**). Domain I (DI, blue), domain II (DII, green), domain III (DIII, yellow), domain IV (DIV, orange), and domain V (DV, red) are shown as ribbons. Insets show a zoomed-in view highlighting DI and DV (FD and Cter for VSV).

## Acknowledgements

This study was supported by the Advanced Research Projects Agency for Health APECx (75N99224R00001 to M.M. and D.V.) National Institute of Allergy and Infectious Diseases (DP1AI158186 and 75N93022C00036 to D.V.), a Pew Biomedical Scholars Award (D.V.), an Investigators in the Pathogenesis of Infectious Disease Awards from the Burroughs Wellcome Fund (D.V.), the University of Washington Arnold and Mabel Beckman cryoEM center and the National Institute of Health grant S10OD032290 (to D.V.). D.V. is an Investigator of the Howard Hughes Medical Institute.

## Author Contributions

M.M. designed the stabilizing mutations and trimers, recombinantly expressed and purified glycoproteins, carried out cryoEM specimen preparation, data collection and processing. M.M. and D.V. built and refined atomic models, analyzed the data, and wrote the manuscript. D.V. supervised the project.

## References

1. Bjornevik, K. et al. Longitudinal analysis reveals high prevalence of Epstein-Barr virus associated with multiple sclerosis. Science 375, 296–301 (2022).

2. Wong, Y., Meehan, M. T., Burrows, S. R., Doolan, D. L. & Miles, J. J. Estimating the global burden of Epstein–Barr virus-related cancers. J. Cancer Res. Clin. Oncol. 148, 31–46 (2022).

3. Damania, B., Kenney, S. C. & Raab-Traub, N. Epstein-Barr virus: Biology and clinical disease. Cell 185, 3652–3670 (2022).

4. Cannon, M. J. Congenital cytomegalovirus (CMV) epidemiology and awareness. J. Clin. Virol. 46 Suppl 4, S6–10 (2009).

5. Itzhaki, R. F. Overwhelming Evidence for a Major Role for Herpes Simplex Virus Type 1 (HSV1) in Alzheimer’s Disease (AD); Underwhelming Evidence against. Vaccines (Basel) 9, (2021).

6. Michalik, F. et al. The effect of herpes zoster vaccination on the occurrence of deaths due to dementia in England and Wales. medRxiv (2023) doi:10.1101/2023.09.08.23295225.

7. Shah, S. et al. Herpes zoster vaccination and the risk of dementia: A systematic review and meta-analysis. Brain Behav. 14, e3415 (2024).

8. Eyting, M., Xie, M., Heß, S., Heß, S. & Geldsetzer, P. Causal evidence that herpes zoster vaccination prevents a proportion of dementia cases. medRxiv (2023) doi:10.1101/2023.05.23.23290253.

9. Smatti, M. K. et al. Epstein-Barr Virus Epidemiology, Serology, and Genetic Variability of LMP-1 Oncogene Among Healthy Population: An Update. Front. Oncol. 8, 211 (2018).

10. Cannon, M. J., Schmid, D. S. & Hyde, T. B. Review of cytomegalovirus seroprevalence and demographic characteristics associated with infection. Rev. Med. Virol. 20, 202–213 (2010).

11. Chemaitelly, H., Nagelkerke, N., Omori, R. & Abu-Raddad, L. J. Characterizing herpes simplex virus type 1 and type 2 seroprevalence declines and epidemiological association in the United States. PLoS One 14, e0214151 (2019).

12. Reynolds, M. A., Kruszon-Moran, D., Jumaan, A., Schmid, D. S. & McQuillan, G. M. Varicella seroprevalence in the U.S.: data from the National Health and Nutrition Examination Survey, 1999-2004. Public Health Rep. 125, 860–869 (2010).

13. Vázquez, M. et al. Effectiveness over time of varicella vaccine. JAMA 291, 851–855 (2004).

14. Oxman, M. N. et al. A vaccine to prevent herpes zoster and postherpetic neuralgia in older adults. N. Engl. J. Med. 352, 2271–2284 (2005).

15. Lopez, A. S., Zhang, J. & Marin, M. Epidemiology of Varicella During the 2-Dose Varicella Vaccination Program - United States, 2005-2014. MMWR Morb. Mortal. Wkly. Rep. 65, 902–905 (2016).

16. Lecrenier, N. et al. Development of adjuvanted recombinant zoster vaccine and its implications for shingles prevention. Expert Rev. Vaccines 17, 619–634 (2018).

17. Tricco, A. C. et al. Efficacy, effectiveness, and safety of herpes zoster vaccines in adults aged 50 and older: systematic review and network meta-analysis. BMJ 363, k4029 (2018).

18. Cunningham, A. L. et al. Efficacy of the Herpes Zoster Subunit Vaccine in Adults 70 Years of Age or Older. N. Engl. J. Med. 375, 1019–1032 (2016).

19. Bender, F. C. et al. Antigenic and mutational analyses of herpes simplex virus glycoprotein B reveal four functional regions. J. Virol. 81, 3827–3841 (2007).

20. Bootz, A. et al. Protective capacity of neutralizing and non-neutralizing antibodies against glycoprotein B of cytomegalovirus. PLoS Pathog. 13, e1006601 (2017).

21. Cairns, T. M. et al. Mechanism of neutralization of herpes simplex virus by antibodies directed at the fusion domain of glycoprotein B. J. Virol. 88, 2677–2689 (2014).

22. Hong, J. et al. Glycoprotein B Antibodies Completely Neutralize EBV Infection of B Cells. Front. Immunol. 13, 920467 (2022).

23. Snijder, J. et al. An Antibody Targeting the Fusion Machinery Neutralizes Dual-Tropic Infection and Defines a Site of Vulnerability on Epstein-Barr Virus. Immunity 48, 799–811.e9 (2018).

24. Sun, C. et al. A gB nanoparticle vaccine elicits a protective neutralizing antibody response against EBV. Cell Host Microbe 31, 1882–1897.e10 (2023).

25. Bu, W. et al. Immunization with Components of the Viral Fusion Apparatus Elicits Antibodies That Neutralize Epstein-Barr Virus in B Cells and Epithelial Cells. Immunity 50, 1305–1316.e6 (2019).

26. Parsons, A. J. et al. Development of broadly neutralizing antibodies targeting the cytomegalovirus subdominant antigen gH. Commun Biol 5, 387 (2022).

27. Zhu, Q.-Y. et al. A potent and protective human neutralizing antibody targeting a novel vulnerable site of Epstein-Barr virus. Nat. Commun. 12, 6624 (2021).

28. Fouts, A. E. et al. Mechanism for neutralizing activity by the anti-CMV gH/gL monoclonal antibody MSL-109. Proc. Natl. Acad. Sci. U. S. A. 111, 8209–8214 (2014).

29. Connolly, S. A., Jardetzky, T. S. & Longnecker, R. The structural basis of herpesvirus entry. Nat. Rev. Microbiol. 19, 110–121 (2021).

30. Chen, W.-H. et al. Epstein-Barr virus gH/gL has multiple sites of vulnerability for virus neutralization and fusion inhibition. Immunity 55, 2135–2148.e6 (2022).

31. Cooper, R. S. & Heldwein, E. E. Herpesvirus gB: A Finely Tuned Fusion Machine. Viruses 7, 6552–6569 (2015).

32. Chandramouli, S. et al. Structure of HCMV glycoprotein B in the postfusion conformation bound to a neutralizing human antibody. Nat. Commun. 6, 8176 (2015).

33. Pötzsch, S. et al. B cell repertoire analysis identifies new antigenic domains on glycoprotein B of human cytomegalovirus which are target of neutralizing antibodies. PLoS Pathog. 7, e1002172 (2011).

34. Zhang, X. et al. Protective anti-gB neutralizing antibodies targeting two vulnerable sites for EBV-cell membrane fusion. Proc. Natl. Acad. Sci. U. S. A. 119, e2202371119 (2022).

35. Oliver, S. L. et al. A glycoprotein B-neutralizing antibody structure at 2.8 Å uncovers a critical domain for herpesvirus fusion initiation. Nat. Commun. 11, 4141 (2020).

36. Sun, C. et al. The structure of HSV-1 gB bound to a potent neutralizing antibody reveals a conservative antigenic domain across herpesviruses. hLife 2, 141–146 (2024).

37. Oliver, S. L. et al. The N-terminus of varicella-zoster virus glycoprotein B has a functional role in fusion. PLoS Pathog. 17, e1008961 (2021).

38. Vollmer, B. et al. The prefusion structure of herpes simplex virus glycoprotein B. Sci Adv 6, (2020).

39. Zhou, M. et al. Targeted mutagenesis of the herpesvirus fusogen central helix captures transition states. Nat. Commun. 14, 7958 (2023).

40. Liu, Y. et al. Prefusion structure of human cytomegalovirus glycoprotein B and structural basis for membrane fusion. Sci Adv 7, (2021).

41. Harrison, S. C. Viral membrane fusion. Nat. Struct. Mol. Biol. 15, 690–698 (2008).

42. Roche, S., Rey, F. A., Gaudin, Y. & Bressanelli, S. Structure of the prefusion form of the vesicular stomatitis virus glycoprotein G. Science 315, 843–848 (2007).

43. White, J. M., Delos, S. E., Brecher, M. & Schornberg, K. Structures and mechanisms of viral membrane fusion proteins: multiple variations on a common theme. Crit. Rev. Biochem. Mol. Biol. 43, 189–219 (2008).

44. Roche, S., Bressanelli, S., Rey, F. A. & Gaudin, Y. Crystal structure of the low-pH form of the vesicular stomatitis virus glycoprotein G. Science 313, 187–191 (2006).

45. Sanders, R. W. & Moore, J. P. Virus vaccines: proteins prefer prolines. Cell Host Microbe 29, 327–333 (2021).

46. Bowen, J. E., et al. SARS-CoV-2 spike conformation determines plasma neutralizing activity elicited by a wide panel of human vaccines. Sci Immunol 7, eadf1421 (2022).

47. McLellan, J. S. et al. Structure-based design of a fusion glycoprotein vaccine for respiratory syncytial virus. Science 342, 592–598 (2013).

48. Sanders, R. W. et al. Stabilization of the soluble, cleaved, trimeric form of the envelope glycoprotein complex of human immunodeficiency virus type 1. J. Virol. 76, 8875–8889 (2002).

49. Pallesen, J. et al. Immunogenicity and structures of a rationally designed prefusion MERS-CoV spike antigen. Proc. Natl. Acad. Sci. U. S. A. 114, E7348–E7357 (2017).

50. Vitu, E., Sharma, S., Stampfer, S. D. & Heldwein, E. E. Extensive mutagenesis of the HSV-1 gB ectodomain reveals remarkable stability of its postfusion form. J. Mol. Biol. 425, 2056–2071 (2013).

51. Silverman, J. L., Sharma, S., Cairns, T. M. & Heldwein, E. E. Fusion-deficient insertion mutants of herpes simplex virus type 1 glycoprotein B adopt the trimeric postfusion conformation. J. Virol. 84, 2001–2012 (2010).

52. Sorem, J. & Longnecker, R. Cleavage of Epstein-Barr virus glycoprotein B is required for full function in cell-cell fusion with both epithelial and B cells. J. Gen. Virol. 90, 591–595 (2009).

53. Dauparas, J. et al. Robust deep learning-based protein sequence design using ProteinMPNN. Science 378, 49–56 (2022).

54. Jumper, J. et al. Highly accurate protein structure prediction with AlphaFold. Nature 596, 583–589 (2021).

55. Walls, A. C. et al. Structure, Function, and Antigenicity of the SARS-CoV-2 Spike Glycoprotein. Cell 181, 281–292.e6 (2020).

56. Yang, F. et al. Structural Analysis of Rabies Virus Glycoprotein Reveals pH-Dependent Conformational Changes and Interactions with a Neutralizing Antibody. Cell Host Microbe 27, 441–453.e7 (2020).

57. Sponholtz, M. R. et al. Structure-based design of a soluble human cytomegalovirus glycoprotein B antigen stabilized in a prefusion-like conformation. bioRxiv 2024.02.10.579772 (2024) doi:10.1101/2024.02.10.579772.

58. Bale, J. B. et al. Accurate design of megadalton-scale two-component icosahedral protein complexes. Science 353, 389–394 (2016).

59. Manley, K. et al. Human cytomegalovirus escapes a naturally occurring neutralizing antibody by incorporating it into assembling virions. Cell Host Microbe 10, 197–209 (2011).

60. Ohlin, M., Sundqvist, V. A., Mach, M., Wahren, B. & Borrebaeck, C. A. Fine specificity of the human immune response to the major neutralization epitopes expressed on cytomegalovirus gp58/116 (gB), as determined with human monoclonal antibodies. J. Virol. 67, 703–710 (1993).

61. Ye, X. et al. Recognition of a highly conserved glycoprotein B epitope by a bivalent antibody neutralizing HCMV at a post-attachment step. PLoS Pathog. 16, e1008736 (2020).

62. Jenks, J. A. et al. A single, improbable B cell receptor mutation confers potent neutralization against cytomegalovirus. PLoS Pathog. 19, e1011107 (2023).

63. Spindler, N. et al. Structural basis for the recognition of human cytomegalovirus glycoprotein B by a neutralizing human antibody. PLoS Pathog. 10, e1004377 (2014).

64. Maginnis, M. S. Virus-Receptor Interactions: The Key to Cellular Invasion. J. Mol. Biol. 430, 2590–2611 (2018).

65. Ma, X. et al. HIV-1 Env trimer opens through an asymmetric intermediate in which individual protomers adopt distinct conformations. Elife 7, (2018).

66. Ozorowski, G. et al. Open and closed structures reveal allostery and pliability in the HIV-1 envelope spike. Nature 547, 360–363 (2017).

67. Lyumkis, D. et al. Cryo-EM structure of a fully glycosylated soluble cleaved HIV-1 envelope trimer. Science 342, 1484–1490 (2013).

68. Julien, J. P. et al. Crystal structure of a soluble cleaved HIV-1 envelope trimer. Science 342, 1477–1483 (2013).

69. Modis, Y., Ogata, S., Clements, D. & Harrison, S. C. Structure of the dengue virus envelope protein after membrane fusion. Nature 427, 313–319 (2004).

70. Rey, F. A., Heinz, F. X., Mandl, C., Kunz, C. & Harrison, S. C. The envelope glycoprotein from tick-borne encephalitis virus at 2 Å resolution. Nature 375, 291–298 (1995).

71. Medits, I. et al. Extensive flavivirus E trimer breathing accompanies stem zippering of the post-fusion hairpin. EMBO Rep. 21, e50069 (2020).

72. Stiasny, K., Kössl, C., Lepault, J., Rey, F. A. & Heinz, F. X. Characterization of a structural intermediate of flavivirus membrane fusion. PLoS Pathog. 3, e20 (2007).

73. Bressanelli, S. et al. Structure of a flavivirus envelope glycoprotein in its low-pH-induced membrane fusion conformation. EMBO J. 23, 728–738 (2004).

74. Kostyuchenko, V. A. et al. Structure of the thermally stable Zika virus. Nature 533, 425–428 (2016).

75. Sirohi, D. et al. The 3.8 Å resolution cryo-EM structure of Zika virus. Science 352, 467–470 (2016).

76. Nicola, A. V. Herpesvirus Entry into Host Cells Mediated by Endosomal Low pH. Traffic 17, 965–975 (2016).

77. Stampfer, S. D., Lou, H., Cohen, G. H., Eisenberg, R. J. & Heldwein, E. E. Structural basis of local, pH-dependent conformational changes in glycoprotein B from herpes simplex virus type 1. J. Virol. 84, 12924–12933 (2010).

78. Qiao, H. et al. Specific single or double proline substitutions in the ‘spring-loaded’ coiled-coil region of the influenza hemagglutinin impair or abolish membrane fusion activity. J. Cell Biol. 141, 1335–1347 (1998).

79. Craig, D. B. & Dombkowski, A. A. Disulfide by Design 2.0: a web-based tool for disulfide engineering in proteins. BMC Bioinformatics 14, 346 (2013).

80. Backovic, M., Longnecker, R. & Jardetzky, T. S. Structure of a trimeric variant of the Epstein-Barr virus glycoprotein B. Proc. Natl. Acad. Sci. U. S. A. 106, 2880–2885 (2009).

81. Watson, J. L. et al. De novo design of protein structure and function with RFdiffusion. Nature 620, 1089–1100 (2023).

82. Suloway, C. et al. Automated molecular microscopy: the new Leginon system. J. Struct. Biol. 151, 41–60 (2005).

83. Punjani, A., Rubinstein, J. L., Fleet, D. J. & Brubaker, M. A. cryoSPARC: algorithms for rapid unsupervised cryo-EM structure determination. Nat. Methods 14, 290–296 (2017).

84. Russo, C. J. & Passmore, L. A. Electron microscopy: Ultrastable gold substrates for electron cryomicroscopy. Science 346, 1377–1380 (2014).

85. Tegunov, D. & Cramer, P. Real-time cryo-electron microscopy data preprocessing with Warp. Nat. Methods 16, 1146–1152 (2019).

86. Bepler, T. et al. Positive-unlabeled convolutional neural networks for particle picking in cryo-electron micrographs. Nat. Methods 16, 1153–1160 (2019).

87. Zivanov, J. et al. New tools for automated high-resolution cryo-EM structure determination in RELION-3. Elife 7, (2018).

88. Zivanov, J., Nakane, T. & Scheres, S. H. W. A Bayesian approach to beam-induced motion correction in cryo-EM single-particle analysis. IUCrJ 6, 5–17 (2019).

89. Punjani, A., Zhang, H. & Fleet, D. J. Non-uniform refinement: adaptive regularization improves single-particle cryo-EM reconstruction. Nat. Methods 17, 1214–1221 (2020).

90. Rosenthal, P. B. & Henderson, R. Optimal determination of particle orientation, absolute hand, and contrast loss in single-particle electron cryomicroscopy. J. Mol. Biol. 333, 721–745 (2003).

91. Chen, S. et al. High-resolution noise substitution to measure overfitting and validate resolution in 3D structure determination by single particle electron cryomicroscopy. Ultramicroscopy 135, 24–35 (2013).

92. Pettersen, E. F. et al. UCSF Chimera--a visualization system for exploratory research and analysis. J. Comput. Chem. 25, 1605–1612 (2004).

93. Nakane, T. & Scheres, S. H. W. Multi-body Refinement of Cryo-EM Images in RELION. Methods Mol. Biol. 2215, 145–160 (2021).

94. He, J., Li, T. & Huang, S.-Y. Improvement of cryo-EM maps by simultaneous local and non-local deep learning. Nat. Commun. 14, 3217 (2023).

95. Emsley, P., Lohkamp, B., Scott, W. G. & Cowtan, K. Features and development of Coot. Acta Crystallogr. D Biol. Crystallogr. 66, 486–501 (2010).

96. Casañal, A., Lohkamp, B. & Emsley, P. Current developments in Coot for macromolecular model building of Electron Cryo-microscopy and Crystallographic Data. Protein Sci. 29, 1069–1078 (2020).

97. Croll, T. I. ISOLDE: a physically realistic environment for model building into low-resolution electron-density maps. Acta Crystallogr D Struct Biol 74, 519–530 (2018).

98. Goddard, T. D. et al. UCSF ChimeraX: Meeting modern challenges in visualization and analysis. Protein Sci. 27, 14–25 (2018).

99. Wang, R. Y. et al. Automated structure refinement of macromolecular assemblies from cryo-EM maps using Rosetta. Elife 5, (2016).

100. Frenz, B. et al. Automatically Fixing Errors in Glycoprotein Structures with Rosetta. Structure 27, 134–139.e3 (2019).

101. Chen, V. B. et al. MolProbity: all-atom structure validation for macromolecular crystallography. Acta Crystallogr. D Biol. Crystallogr. 66, 12–21 (2010).

102. Barad, B. A. et al. EMRinger: side chain-directed model and map validation for 3D cryo-electron microscopy. Nat. Methods 12, 943–946 (2015).

103. Liebschner, D. et al. Macromolecular structure determination using X-rays, neutrons and electrons: recent developments in Phenix. Acta Crystallogr D Struct Biol 75, 861–877 (2019).

104. Abramson, J. et al. Accurate structure prediction of biomolecular interactions with AlphaFold 3. Nature (2024) doi:10.1038/s41586-024-07487-w.

